# Rewiring Catalytic Craters: A Path for Engineering β-Glucosidases to Improve Glucose Tolerance

**DOI:** 10.1101/2025.01.02.631061

**Authors:** Abhishek B. Suryawanshi, Rajiv K. Bedi, Meera Pawar, Santosh Noronha, Prasenjit Bhaumik

**Author notes:** Prasenjit Bhaumik, Department of Biosciences and Bioengineering, Indian Institute of Technology Bombay, Powai, Mumbai-400076, India.

## Abstract

Bioethanol, a sustainable alternative to fossil fuels, is produced from cellulose via enzymatic saccharification. β-Glucosidase, a key enzyme in this process, hydrolyses disaccharides into glucose but is limited by feedback inhibition. This study investigated GH1 β-glucosidase from a soil metagenome (UnBGl1) to understand and overcome this limitation. We solved the near-atomic resolution crystal structure of UnBGl1 at 1.15 Å as an apo enzyme and report the first high-resolution crystal structures of the enzyme in its pre-hydrolytic state as a cellobiose complex and covalent intermediate-bound state, capturing key stages of its catalytic mechanism. Structural analysis revealed three crucial glucose-binding subsites in UnBGl1’s crater. Glucose binding at the −1 subsite induced a 1.4 Å shift in E370, increasing the distance between catalytic glutamates and reducing enzymatic activity. The C188V variant, generated by rational engineering, significantly improved glucose tolerance to 2.5 M. Additionally, the H261W mutation at the +2 subsite enhanced kinetic properties by improving cellobiose affinity (*K*_m_ = 22.87 ± 1.1 mM) and shifted the optimal pH to 5.5 from 6.0. Comparative structural analysis with other glucose-tolerant GH1 β-glucosidases revealed conserved residues at the −1 subsite crucial for substrate stabilization and +1 subsite residues interacting with glucose, offering targets for further optimization. Engineered UnBGl1 variants retained high stability and activity on sugarcane bagasse, demonstrating their potential for industrial cellulase cocktails. These findings provide a robust framework for engineering β-glucosidases with enhanced glucose tolerance and catalytic efficiency, paving the way for improved bioethanol production and contributing to sustainable energy solutions.

## Introduction

The dual threat of climate change and finite fossil fuel reserves have led us to search and development of environment-friendly, renewable, carbon-neutral energy solutions. Bioethanol is one such energy source, which is derived from plant-based lignocellulose. The main benefit of employing second-generation bioethanol is that it requires the lignocellulosic biomass as primary resource, which avoids interference with food crops and can readily be grown on poor-quality marginal land with little water and fertilizers^1^. However, the presence of lignin and hemicellulose can hinder the process of bioethanol production. Therefore, the separation of lignin and hemicellulose from the lignocellulose is done by several physical, physicochemical, chemical and biological approaches to obtain cellulose^2^. Cellulose is the most abundant and renewable carbohydrate bio-polymer comprising of β-1,4-linked glucose monomers^3^. Hydrolysis of cellulose by cellulases produces glucose to produce ethanol. Cellulases are enzymes complexes, including endo-glucanases, cellobiohydrolases, exo-glucanases and β-glucosidases. These enzymes synergistically hydrolyse cellulose into fermentable sugars^4–6^. Endo-glucanase acts randomly on the amorphous area of the cellulose and generates various lengths of oligosaccharides; cellobiohydrolases and exo-glucanases catalyze the release of glucose or cellobiose from the reducing and non-reducing end of polysaccharide and finally β-glucosidase hydrolyses and produces glucose from released cellobiose^5,7^.

Cellobiose acts as an inhibitor for endo-glucanase and exo-glucanase^4^. β-glucosidase, the final enzyme in the saccharification process, catalyses the hydrolysis of cellobiose and other smaller oligosaccharides. Even if β-glucosidase can reduce the inhibitory effect of cellobiose, it produces glucose, which at higher concentrations acts as an inhibitor for β-glucosidase^8,9^. Increased concentration of the sugars produced during the saccharification process in the reactors is responsible for the increase in viscosity and raises the difficulty of the product diffusion out of the active site of β-glucosidase^10^. At high concentrations, glucose can either block the β-glucosidase active site for substrate entry or prevent product exit^11,12^. Moreover, it is worth mentioning that there is a correlation between glucose tolerance and a stimulating effect triggered by glucose in β-glucosidases. This correlation only applies to some members of the glycoside hydrolase family 1 (GH1) β-glucosidases^12^. Stimulation effect in β-glucosidases can be observed due to the binding of glucose to the secondary site, whereas the enzyme gets inhibited when glucose binds and blocks the active site^13^. A deeper understanding of feedback inhibition of β-glucosidases by glucose is highly essential and would contribute to the development of efficient enzymes for the industrial saccharification process.

Bioethanol producing industries require the β-glucosidases with higher activity and lower *K*_m_ for saccharification to reduce the burden of inhibition exerted by cellobiose^14^ as insufficient activity leads to build-up of cellobiose during cellulose hydrolysis^15,16^. One of the strategies used by industry to avoid feedback inhibition is the implication of simultaneous saccharification and fermentation (SSF) method for bioethanol production where both saccharification and fermentation are performed in a single reactor. Hence, the released glucose from the saccharification process is used immediately for bioethanol production^17^. Generally, SSF occurs at lower temperatures as organisms like yeast that ferment sugar prefers lower temperatures. Therefore, it is essential to develop enzymes that do not undergo product inhibition and have higher activity and stability at lower temperatures. Until now, no proper comprehensive studies have been conducted that reveal the molecular basis of glucose tolerance in β-glucosidase^18^. Here we present extensive structural and biochemical studies to understand the mechanistic details of product inhibition and designing strategies for improving the glucose tolerance of GH1 β-glucosidases.

β-glucosidase (EC 3.2.1.21) is an enzyme that hydrolyses the compounds with β-D-glycosidic linkage^18,19^. Several studies have reported functional characterizations of this enzyme due to its universal distribution and well-defined variety of substrates^20^. β-glucosidase has been classified into different glycoside hydrolase families: GH1, GH3, GH5, GH9 and GH30. Among them, the GH1 family contains the most significant number of characterized β-glucosidases with the classic (α/β)_8_ TIM-barrel structure^21,22^. GH1 family β-glucosidases are enzymes which catalyse hydrolysis of β-D-glycosidic bonds by retaining a double displacement mechanism^23^. The conserved carboxylic acid residues present in the β-strands 4 and 7 of the enzyme serve as both the general acid-base catalyst and the nucleophile during the catalytic reaction^24^. The catalytic acid/base is located on the TXNEP motif at the end of strand 4 (where X is a hydrophobic amino acid residue), and the catalytic nucleophile is located on the I/VTENG motif at the end of strand 7^25^. The catalysis of β-glucosidase has been studied by many methods, including pH-activity analysis, mutagenesis, suicide inactivator, isotope effects, interacting studies with alternative substrates or inhibitors and 3-Dimensional (3D) structural visualization. However, the coordinated conformational changes enabling substrate distortion and glycosidic bond cleavage have remained primarily obscured over decades of study^8,26^. Extensive structural understanding of substrate hydrolysis is still poorly understood; therefore, structural studies with native GH1 enzymes are necessary.

In this study, we have performed biochemical characterizations of a GH1 enzyme, UnBGl1, a β-glucosidase identified from soil metagenome^27^; and solved multiple high-resolution crystal structures of native and variants of this enzyme. Further, through our crystallographic studies, we have also demonstrated three stages of cellobiose hydrolysis catalytic mechanism showing the substrate (cellobiose), covalently linked glycosyl reaction intermediate and product (glucose) binding at the catalytic centre of UnBGl1. Our study is the first report that captured the covalently bound reaction intermediate to the catalytic site of a GH1 β-glucosidase. Apo, ligand (glucose, cellobiose, thio-glucose) complexed and glucose-bound covalent intermediate structures aided in identifying the conserved product binding sites in UnBGl1. Structural analysis has provided insights into understanding the molecular basis to increase glucose tolerance and improve the kinetic properties of UnBGl1. These findings can be applied to other GH1 family β-glucosidase to enhance their glucose tolerance.

## Results

### Kinetic and biochemical properties of native UnBGl1

The C-terminal His_6_ tagged recombinant UnBGl1 was overexpressed as a soluble protein by induction of 0.4 mM IPTG at 24 ℃ for 16 hrs. After the purification by Ni-NTA affinity and size exclusion chromatography, highly pure UnBGl1 was obtained. The chromatogram (Figure 1a) showed the single elution peak, indicating the purity of the protein, which was confirmed by SDS-PAGE. Highly purified UnBGl1 was further used for biochemical and kinetic characterizations and crystallization.

**Figure 1:**
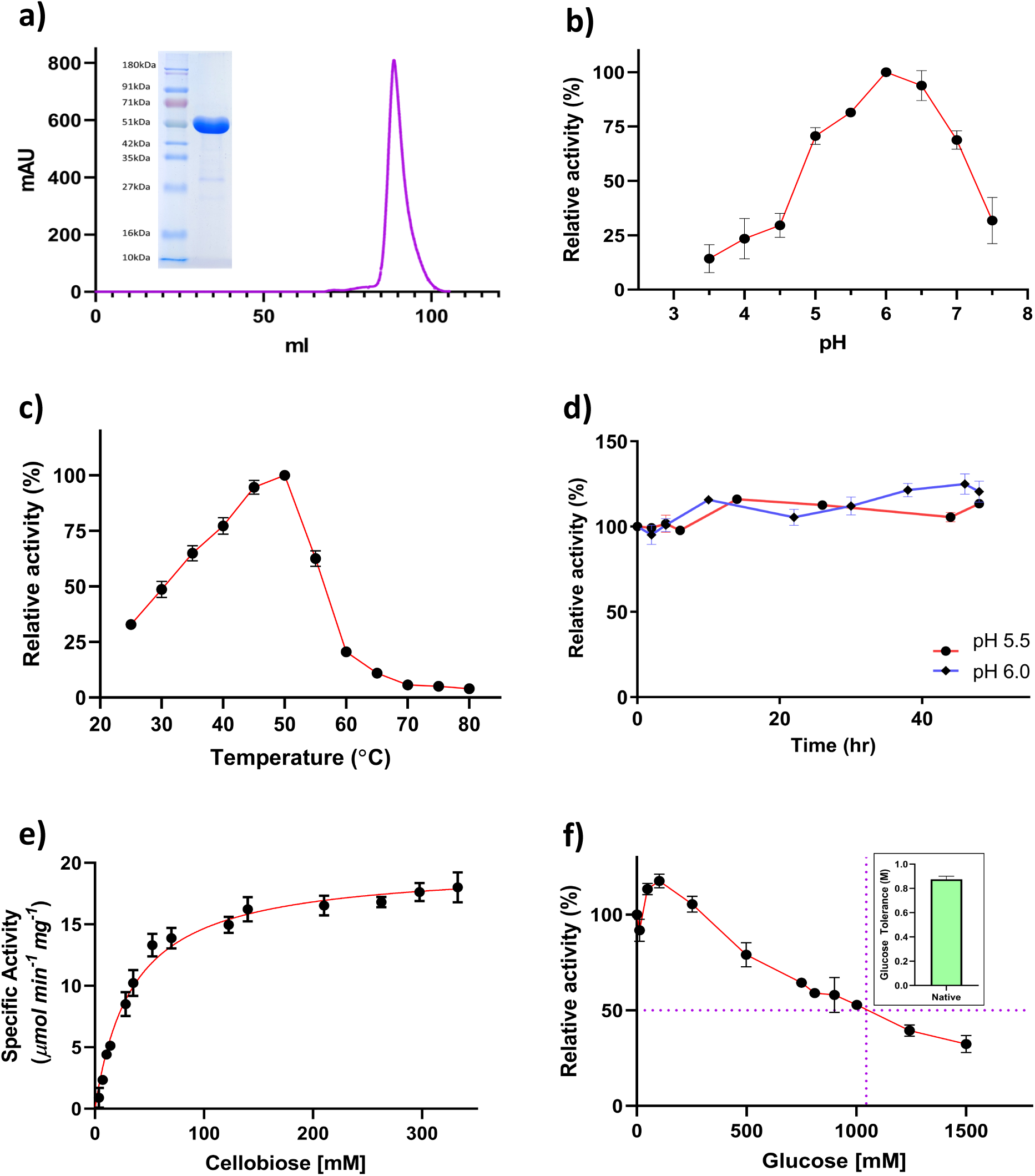
Biochemical properties of native UnBGl1. **a)** Size exclusion chromatogram (purple colour line) indicating the elution of purified native UnBGl1 from Superdex 200 16/600 column. Single band near 51 kDa on SDS-PAGE (inset) represents the purity of the enzyme. **b, c)** Graphs illustrate the optimum pH of 6.0 and temperature of 50 °C for cellobiose hydrolysis, respectively. Data points are shown in black solid circles and connected with the red line. **d)** stability of native UnBGl1 at pH 5.5 (red) and pH 6.0 (purple) at 30°C. **e)** Kinetics profile of native UnBGl1 showing the substrate saturation for cellobiose hydrolysis. **f)** The graph depicts UnBGl1 glucose tolerance level expressed as concentration of glucose that causes loss of 50 % enzyme activity. Inset is a bar graph representation of glucose tolerance. All experiments were performed in triplicates (n = 3) with error bar representing ± SEM.

We performed 4-nitrophenyl β-D-glucopyranoside (pNPG) and cellobiose hydrolysis reactions at various pHs and temperatures to determine the best conditions for the catalytic reaction by UnBGl1. Native UnBGl1 activity was investigated throughout a pH range of 3.5 - 8.0, with reactions occurring at 55 °C for 10 minutes. At pH 6.0, the highest activity was observed for pNPG (Supplementary Figure S1a) and cellobiose hydrolysis (Figure 1b). In a pH range of 5.5 - 7.0, the enzyme showed more than 70 % of its highest activity. Below pH 5.0 and above pH 7.5, activity was less than 50 % of its maximum value. Native UnBGl1 showed increased activity in the temperature range of 45 °C - 55 °C, with maximum activity at 55 °C for pNPG hydrolysis (Supplementary Figure S1b) and 50 °C for cellobiose hydrolysis (Figure 1c). The enzyme activity dropped to less than 30 % at temperatures above 55 °C. Purified native UnBGl1 is stable at pH 5.5 and pH 6.0 for 48 hrs. and retains almost 100% of its activity after 48 hrs. (Figure 1d).

The kinetic properties of native UnBGl1 were initially determined with its artificial substrate pNPG, which, upon hydrolysis, forms p-nitrophenol (pNP), which is a yellow colour in the alkaline pH. The substrate saturation assays were performed to determine the kinetic parameters (*K*_m_*, V_m_*_ax_) of UnBGl1. The native enzyme followed Michaelis-Menten kinetics, with *V*_max_ value of 36.14 ± 0.50 µmol min^-1^mg^-1^, *K*_m_ value of 0.92 ± 0.05 mM and catalytic efficiency (*k*_cat_*/ K*_m_) value of 34.36 ± 11 mM^-1^ s^-1^ (Supplementary Figure S1c). We also determined the kinetic parameters of native UnBGl1 for hydrolysis of cellobiose, a natural substrate of β-glucosidase. For cellobiose hydrolysis, UnBGl1 attained the *V_max_* of 19.74 ± 0.37 µmol min^-1^mg^-1^ with *K*_m_ of 34.61 ± 2.4 mM and *k*_cat_*/ K*_m_ of 0.47 ± 0.17l mM^-1^ s^-1^ (Figure 1e). The glucose tolerance level of the native UnBGl1 was determined. It was observed that native UnBGl1 loses its 50 % activity in the presence of around 1.0 M (Figure 1f) glucose concentration.

### Substrate, product and reaction intermediate binding in the native UnBGl1

The near atomic-resolution crystal structure of the native UnBGl1 was solved as apo-enzyme at 1.15 Å (PDB ID. 9JLZ) (Table 1: part I). The structure has been well refined with an acceptable range of *R*_work_ and *R*_free_. This 469 residue-containing enzyme has a spherical shape. The overall dimension of UnBGl1 structure is approximately 57.5 × 50.5 × 68.3 Å. This enzyme belongs to the GH1 family and has a well-characterized typical (α/β)_8_ TIM-barrel fold (Figure 2a, Supplementary movie S1). The structure of UnBGl1 was compared with the structure of its close well-characterized homologous (50.8 % sequence identity) (Supplementary Figure S2a) β-glucosidase from *Humicola insolens* (HiBG) (PDB ID. 4MDO)^13^ and root mean square deviation (r.m.s.d.) value was 0.695 Å. (Supplementary Figure S2b). Structural alignment of these two enzymes helped to identify the catalytic crater and some gatekeeper residues of UnBGl1. E185 (acid-base) and E370 (nucleophile) are the catalytic residues (Figure 2b). H140, N184, and W417 are the supporting active site residues, and E424, W343, F313, C188, L187, Y312, I192, and H261 (Supplementary Figure S2c) are residues that form the crater of the UnBGl1. The crater of apo UnBGl1 is filled with water molecules. In the catalytic crater of the apo enzyme structure, we also observed two glycerol molecules which might have appeared from the cryo-protectant solution (Supplementary Figure S2d).

**Table 1.** part I: Data collection and refinement statistics of UnBGl1 variants.

| **Data collection statistics<sup>a</sup>** | **UnBGI1 Apo** | **UnBGI1 Glucose** | **UnBGI1 Cellobiose** | **UnBGI1 Covalent Intermediate** | **UnBGI1 Thio-glucose** | **UnBGI1 C188V Apo** |
| --- | --- | --- | --- | --- | --- | --- |
| Wavelength (Å) | 0.97893 | 1.5418 | 1.5418 | 1.5418 | 1.5418 | 0.97949 |
| Space group | *P*2<sub>1</sub>2<sub>1</sub>2<sub>1</sub> | *P*2<sub>1</sub>2<sub>1</sub>2<sub>1</sub> | *P*2<sub>1</sub>2<sub>1</sub>2<sub>1</sub> | *P*2<sub>1</sub>2<sub>1</sub>2<sub>1</sub> | *P*2<sub>1</sub>2<sub>1</sub>2<sub>1</sub> | *P*2<sub>1</sub>2<sub>1</sub>2<sub>1</sub> |
| Unit cell constants |  |  |  |  |  |  |
| *a*, *b*, *c* (Å) | 51.84, 65.99, 138.18 | 51.91, 66.18, 138.17 | 51.47, 66.51, 138.20 | 51.20, 65.80, 137.90 | 51.15, 65.71, 137.66 | 51.88, 66.27, 138.25 |
| $\alpha$, $\beta$, $\gamma$ (°) | 90.0, 90.0, 90.0 | 90.0, 90.0, 90.0 | 90.0, 90.0, 90.0 | 90.0, 90.0, 90.0 | 90.0, 90.0, 90.0 | 90.0, 90.0, 90.0 |
| Resolution (Å) | 40.0-1.15 (1.25-1.15) | 40.0-1.73 (1.80-1.73) | 40.0-1.73 (1.8-1.73) | 30.0-2.0 (2.1-2.0) | 40.0-1.73 (1.83-1.73) | 30.0-1.3 (1.4-1.3) |
| No. of reflections | 1161293 (217836) | 349373 (32373) | 343914 (30879) | 225378 (29989) | 325795 (42715) | 549033 (97401) |
| Unique reflections | 167030 (35537) | 50247 (5305) | 50020 (5356) | 31988 (4234) | 49212 (7451) | 116123 (23019) |
| Completeness (%) | 99.1 (96.3) | 99.4 (94.9) | 99.3 (96.2) | 99.1 (98.1) | 99.9 (99.2) | 98.6 (99.2) |
| Multiplicity | 6.95 (6.12) | 6.95 (6.10) | 6.87 (5.76) | 7.51 (7.08) | 6.62 (5.73) | 4.72 (4.2) |
| *R*<sub>merge</sub> (%) | 7.4 (55.4) | 2.5 (7.0) | 4.8 (45.8) | 30.3 (90.3) | 3.5 (16.7) | 5.1 (43.9) |
| Mean *I*/$\sigma$<*I*> | 13.61 (3.17) | 56.01 (23.33) | 22.91 (3.5) | 9.54 (2.80) | 35.02 (9.99) | 16.62 (3.4) |
| CC<sub>1/2</sub> (%) | 99.9 (85.2) | 100 (99.7) | 99.9 (90.9) | 95.7 (76.8) | 100.0 (98.1) | 99.9 (88.0) |
| **Refinement statistics** |  |  |  |  |  |  |
| Resolution (Å) | 39.10-1.15 | 37.8-1.73 | 37.87-1.73 | 29.69-2.0 | 38.73-1.73 | 28.76-1.30 |
| Working set: no. of reflections | 158699 | 47733 | 47521 | 30390 | 46751 | 110314 |
| *R*<sub>factor</sub> (%) | 12.58 | 12.82 | 16.36 | 18.56 | 14.69 | 16.80 |
| Test set: no. of reflections | 8353 | 2513 | 2502 | 1600 | 2461 | 5806 |
| *R*<sub>free</sub> (%) | 14.70 | 15.57 | 20.04 | 23.15 | 18.06 | 18.77 |
| **Number of atoms** |  |  |  |  |  |  |
| Protein atoms | 3728 | 3638 | 3625 | 3584 | 3601 | 3660 |
| Water molecules | 636 | 558 | 381 | 224 | 524 | 593 |
| **Ligands** |  |  |  |  |  |  |
| Glucose | 0 | 3 | 1 | 3 | 1 | 0 |
| Cellobiose | 0 | 0 | 1 | 0 | 0 | 0 |
| Bond lengths (Å) | 0.012 | 0.012 | 0.008 | 0.006 | 0.011 | 0.012 |
| Bond angles (°) | 1.849 | 1.903 | 1.596 | 1.541 | 1.881 | 1.914 |
| **Ramachandran Plot<sup>c</sup> (%)** |  |  |  |  |  |  |
| Favoured region | 96.9 | 96.9 | 96.5 | 96.9 | 97.1 | 96.9 |
| Allowed region | 3.1 | 3.1 | 3.5 | 3.1 | 2.9 | 3.1 |
| Disallowed region | 0 | 0 | 0 | 0 | 0 | 0 |
| PDB code | **9JLZ** | **9J3W** | **9J3Y** | **9J4Q** | **9J4B** | **9J4I** |

<sup>a</sup>Values in parentheses indicate statistics in highest resolution shell.
<sup>b</sup> $R_{\text{merge}} = \sum |I - \langle I \rangle| / \sum I$ where *I* is the observed integrated intensity, $\langle I \rangle$ is the average integrated intensity obtained from multiple measurements, and the summation is overall observed reflections.
<sup>c</sup>Values are obtained from PROCHECK analysis.

**Table 1.** part II: Data collection and refinement statistics of UnBGl1 variants.

| *Data collection statistics<sup>a</sup>* | UnBGI1 C188V Glucose | UnBGI1 H261W Apo | UnBGI1 H261W Glucose | UnBGI1 H261W Cellobiose | UnBGI1 C188V_ H261W Apo | UnBGI1 C188V_ H261W Glucose |
| --- | --- | --- | --- | --- | --- | --- |
| Wavelength (Å) | 1.5418 | 0.97777 | 1.5418 | 0.97777 | 0.97777 | 0.97777 |
| Space group | *P*2<sub>1</sub>2<sub>1</sub>2<sub>1</sub> | *P*2<sub>1</sub>2<sub>1</sub>2<sub>1</sub> | *P*2<sub>1</sub>2<sub>1</sub>2<sub>1</sub> | *P*2<sub>1</sub>2<sub>1</sub>2<sub>1</sub> | *P*2<sub>1</sub>2<sub>1</sub>2<sub>1</sub> | *P*2<sub>1</sub>2<sub>1</sub>2<sub>1</sub> |
| Unit cell constants |  |  |  |  |  |  |
| *a*, *b*, *c* (Å) | 52.11, 66.98, 138.64 | 51.66, 66.14, 137.33 | 51.47, 65.87, 137.35 | 50.53, 65.36, 136.15 | 51.75, 66.38, 137.49 | 52.00, 68.10, 136.00 |
| $\alpha$, $\beta$, $\gamma$ (°) | 90.0, 90.0, 90.0 | 90.0, 90.0, 90.0 | 90.0, 90.0, 90.0 | 90.0, 90.0, 90.0 | 90.0, 90.0, 90.0 | 90.0, 90.0, 90.0 |
| Resolution (Å) | 40.0-1.75 (1.85-1.75) | 40.0-1.8 (1.9-1.8) | 65.87-1.78 (1.81-1.78) | 40.58-1.70 (1.73-1.70) | 30.0-1.55 (1.65-1.55) | 30.0-2.0 (2.1-2.0) |
| No. of reflections | 339608 (48264) | 324512 (48061) | 323344 (14972) | 372926 (18249) | 515477 (72060) | 239076 (32459) |
| Unique reflections | 48288 (7023) | 44406 (6535) | 45575 (2183) | 50499 (2496) | 69348 (11661) | 33409 (4463) |
| Completeness (%) | 96.9 (93.0) | 99.9 (99.8) | 100.0 (99.0) | 100 (99.8) | 99.7 (99.3) | 99.9 (100) |
| Multiplicity | 7.03 (6.87) | 6.75 (7.35) | 7.09 (6.85) | 7.4 (7.3) | 7.43 (6.17) | 7.1 (7.2) |
| *R*<sub>merge</sub> (%) | 7.1 (28.7) | 24.9 (101.1) | 23.9 (78.2) | 14.7 (70.3) | 10.0 (82.7) | 22.6 (87.1) |
| Mean *I*/$\sigma$<*I*> | 19.31 (6.64) | 9.38 (2.93) | 8.8 (2.7) | 12.2 (2.3) | 14.42 (2.45) | 7.44 (2.30) |
| CC<sub>1/2</sub> (%) | 99.8 (95.3) | 99.2 (79.9) | 98.7 (30.8) | 99.3 (82.0) | 99.8 (76.7) | 99.0 (76.0) |
| Refinement statistics |  |  |  |  |  |  |
| Resolution (Å) | 38.04-1.75 | 39.03-1.8 | 59.34-1.78 | 40.58-1.70 | 29.89-1.55 | 28.49-2.0 |
| Working set: no. of reflections | 45873 | 42185 | 43195 | 47867 | 65880 | 31738 |
| *R*<sub>factor</sub> (%) | 15.37 | 13.96 | 17.4 | 14.16 | 13.88 | 16.17 |
| Test set: no. of reflections | 2415 | 2221 | 2309 | 2561 | 3468 | 1671 |
| *R*<sub>free</sub> (%) | 19.11 | 16.78 | 20.74 | 18.00 | 16.30 | 21.7 |
| Number of atoms |  |  |  |  |  |  |
| Protein atoms | 3614 | 3634 | 3701 | 3662 | 3742 | 3580 |
| Water molecules | 574 | 518 | 41 | 410 | 523 | 338 |
| Ligands |  |  |  |  |  |  |
| Glucose | 2 | 0 | 2 | 1 | 0 | 1 |
| Cellobiose | 0 | 0 | 0 | 2 | 0 | 0 |
| Bond lengths (Å) | 0.01 | 0.009 | 0.008 | 0.009 | 0.011 | 0.015 |
| Bond angles (°) | 1.602 | 1.58 | 1.505 | 1.652 | 1.679 | 2.078 |
| Ramachandran Plot<sup>c</sup> (%) |  |  |  |  |  |  |
| Favoured region | 96.5 | 97.4 | 97.2 | 97.2 | 97.0 | 97.3 |
| Allowed region | 3.5 | 2.6 | 2.8 | 2.8 | 3.0 | 2.7 |
| Disallowed region | 0 | 0 | 0 | 0 | 0 | 0 |
| PDB code | **9J42** | **9J43** | **9J44** | **9J45** | **9J49** | **9J4A** |

<sup>a</sup>Values in parentheses indicate statistics in highest resolution shell.
<sup>b</sup> $R_{\text{merge}} = \sum |I - \langle I \rangle| / \sum I$ where *I* is the observed integrated intensity, <*I*> is the average integrated intensity obtained from multiple measurements, and the summation is overall observed reflections.
<sup>c</sup>Values are obtained from PROCHECK analysis.

**Figure 2:**
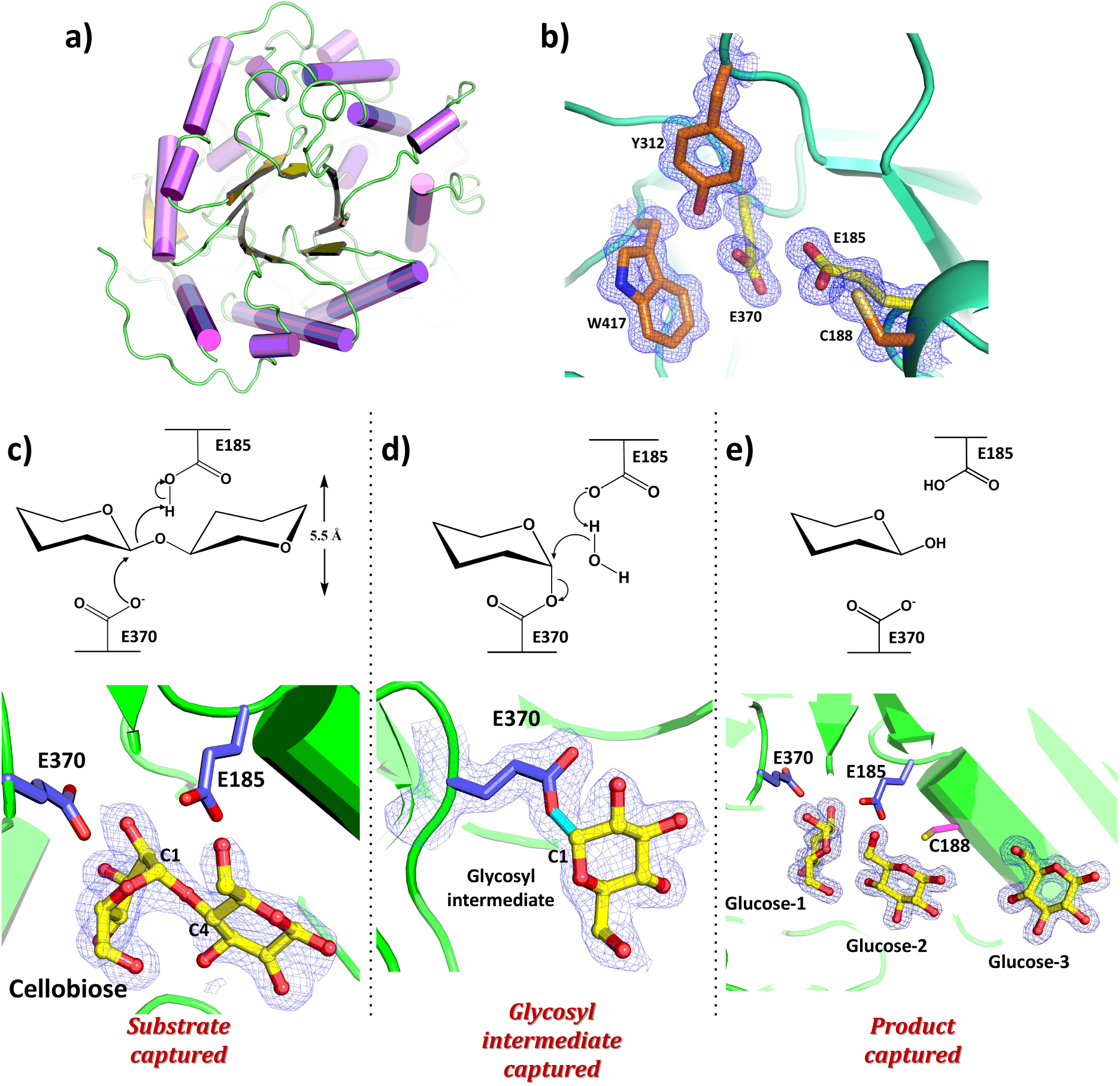
Structural fold, active site architecture and cellobiose hydrolysis mechanism of UnBGl1. **a)** Overall structure of UnBGl1 with (α/β)_8_ TIM-barrel fold is represented as cartoon. Helixes are shown in purple whereas sheets are shown in yellow colour. **b)** The 2*F_o_ - F_c_* map (blue mesh) contoured at 1σ level showing the fully satisfied electron density around active site residues. Catalytic glutamates are presented as yellow stick and other active site residues are represented as orange sticks. **Mechanism of cellobiose hydrolysis captured in the active site of UnBGl1. The catalytic glutamates are shown in purple stick, cellobiose and glucose are represented as ball (red) and stick (yellow) model**. **c)** The 2*F_o_ - F_c_* map contoured at 1 σ level representing the electron density (slate mesh) of cellobiose near the catalytic glutamates suggesting that substrate has been captured just before nucleophilic attack. **d)** The 2*F_o_ - F_c_* map contoured at 1 σ level illustrating the electron density (slate mesh) of glycosyl intermediate covalently linked to catalytic glutamate (E370). **e)** The 2*F_o_ - F_c_* map contoured at 1 σ level showing the electron density (slate mesh) of three glucose molecules at the active site crater of UnBGl1.

Understanding the hydrolysis mechanism of any enzyme in its native state is challenging. However, we aimed to visualize cellobiose hydrolysis by UnBGl1 and understand the molecular details of substrate, intermediate and product binding in this GH1 enzyme active site. To demonstrate the enzymatic activity of UnBGl1 in the crystal state, the crystals were soaked with pNPG substrate. Within a short time of incubation, the surrounding solution started turning yellow (Supplementary movie S2) due to the formation of a yellow product pNP upon hydrolysis of pNPG by UnBGl1 present in the crystal. These observations confirmed that UnBGl1 was enzymatically active in the crystal form and could hydrolyse its substrate cellobiose to glucose. We performed various soaking experiments using native UnBGl1 crystals with multiple concentrations of cellobiose and varied soaking periods to capture substrate, reaction intermediate and glucose in the enzyme’s active site. Cellobiose complexed structure (PDB ID. 9J3Y) was obtained from a short soaking experiment of native UnBGl1 crystals in cellobiose. Upon solving the structure, we found out that the UnBGl1 active site has the substrate cellobiose (Supplementary Figures S3a, S3b and S3c) and the hydrolysis product glucose molecule (Supplementary Figures S3b and S3c). The cellobiose molecule is bound near the catalytic glutamates (E185, E370) (Figure 2c, Supplementary Figure S3a, Supplementary movie S3). At the catalytic crater, aromatic residues like W141, W343, F313, W417 and Y312 dominate the interaction with glucose units of the substrate. In the process of capturing the substrate-bound state of the UnBGl1, we also captured one of the intermediate states of the enzyme reaction. We determined a crystal structure of UnBGl1 complexed to the reaction intermediate at 2.0 Å resolution (PDB ID. 9J4Q) (Table 1: part I). In this structure, we observed that nucleophilic glutamate (E370) formed a covalent bond with the reaction intermediate, which was produced by breaking of β-1,4 glycosidic linkage (Figure 2d, Supplementary Figure S3d) of cellobiose by UnBGl1. We could also see the product glucose in the crater and the reaction intermediate formation during the hydrolysis. Capturing the reaction intermediate in the native and active state of the enzyme through crystallography is one of the complex tasks. Until now, there is no report of capturing the glycosyl intermediate through crystallography using an active β-glucosidase. Our study is the first report to present a high-resolution structure of a covalently bound glycosyl intermediate formed by the action of active native UnBGl1 on cellobiose.

β-glucosidases get inhibited by its product glucose due to the feedback inhibition. We solved the 1.73 Å resolution UnBGl1 glucose complexed structure (PDB ID. 9J3W) (Table 1: part I) that provided insights into crucial residues which facilitate feedback inhibition by the product. Three glucose molecules were discovered during the refinement of the structure (Figure 2e, Supplementary Figure S3e). The glucose bound and cellobiose bound structures did not show any significant difference to the apo structure of native UnBGl1 and have root mean square deviation (r.m.s.d.) of 0.136 Å and 0.148 Å, respectively (Supplementary Figure S3f), based on the overall structural superposition.

For the GH1 family β-glucosidases that follow a retaining reaction mechanism, the typical separation (measured by the distance between CD atoms) of the two carboxylate groups of the catalytic glutamates is ̴ 5.5 Å^23,28^. In the apo UnBGl1 structure, this distance is 5.1 Å (Figure 3a). However, at high glucose concentrations, the sugar molecules bind to all the conserved glucose binding sites. Notably, the carboxylate group of the nucleophilic glutamate (E370) showed 1.4 Å displacement from its original position that is observed for apo or cellobiose bound UnBGl1 (Figure 3b). In the glucose complexed UnBGl1 structure, it is observed that the nucleophilic glutamate (E370) side chain is a bit displaced from its position, and the distance between two catalytic glutamates (E185, E370) increased to 6.0 Å (Figure 3c). Such wide distance between two catalytic glutamates is not optimal to facilitate catalysis. The glucose molecules are stabilized by multiple interactions in the active site crater of UnBGl1.

**Figure 3:**
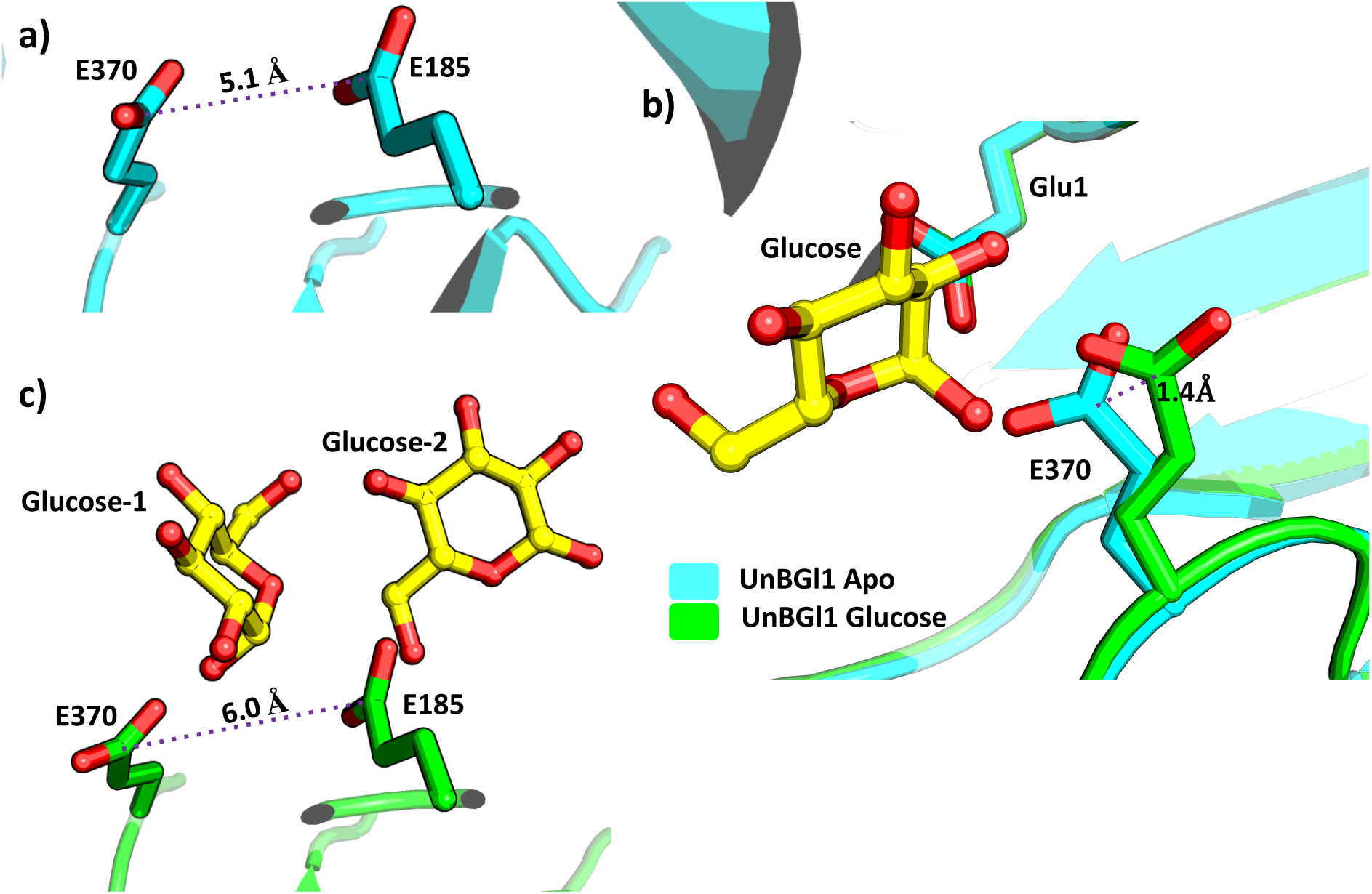
Structural difference at the active site of apo and glucose bound native UnBGl1. The structure of native apo UnBGl1 (cyan) is compared with the glucose complexed structure (green). **a)** Distance between the CD atoms of catalytic glutamates of apo UnBGl1 is shown in dotted line. **b)** Catalytic glutamate (E370) showing 1.4 Å displacement from its original place due to binding of glucose. **c)** Distance between the CD atoms of catalytic glutamates of glucose complexed UnBGl1 is shown in dotted line.

Our data present the first report of successfully captured three distinct stages of cellobiose hydrolysis reaction: (a) substrate (cellobiose) at the catalytic site (Figure 2c), (b) covalently bound glycosyl intermediate to the catalytic glutamate (E370) (Figure 2d) and (c) product (glucose) in the active site crater (Figure 2e) of an active GH1 enzyme, UnBGl1. These high-resolution crystal structures of UnBGl1 provided quite detailed insights into the substrate binding and the conserved glucose binding sites.

### Identification of conserved glucose binding sites in UnBGl1 and rational engineering

From the analysis of the glucose-complexed native UnBGl1 structures, we identified the specific sites where glucose molecules bind in the catalytic crater of this enzyme. These binding sites are designated as −1 subsite, +1 subsite, and +2 subsite (Figure 4a). The cellobiose binds near the catalytic glutamates where the substrate hydrolysis occurs (Figure 2c, Supplementary Figure S3a), and this site is referred as the −1 subsite. Also, in a glucose complexed structure, one glucose molecule is seen at −1 subsite, that contains catalytic residues. Any structural hindrance at −1 subsite may adversely affect the enzyme’s activity, so it is better not to make any structural perturbation at −1 subsite. A second glucose molecule was found at +1 subsite located at the middle of the crater. The glucose molecule is well stabilized at the +1 subsite through interaction with the side chains of residues N241, E185, C188, I192, W343, F313. To understand the importance of residues involved in stabilizing the substrate at both −1 and +1 subsite, a thio-cellobiose soaking experiment with native UnBGl1 crystal was performed. Thio-cellobiose acts as a moderate competitive inhibitor of β-glucosidase and has a similar structural resemblance to its substrate cellobiose^22,29^. To our surprise, no ligand was observed near the catalytic glutamates in the active site of this structure. Instead, a thio-glycosidic enzyme intermediate covalently bound to 188^th^ cysteine (C188) residue at +1 subsite was formed due to which the substrate entry channel is blocked (PDB ID. 9J4B) (Figure 4b, Supplementary Figure S4a). Thio-glucose bound structure suggests possible importance of C188 facilitating feedback inhibition of UnBGl1. We also hypothesized, introducing the hydrophobic interactions in the crater may restrict the entry of glucose to the active site^5^. Therefore, to test our hypothesis cysteine at 188^th^ position at +1 subsite was mutated to valine (C188V) (Supplementary Figure S4b) to increase hydrophobic interactions and further biochemical as well as structural studies of the variant were performed.

**Figure 4:**
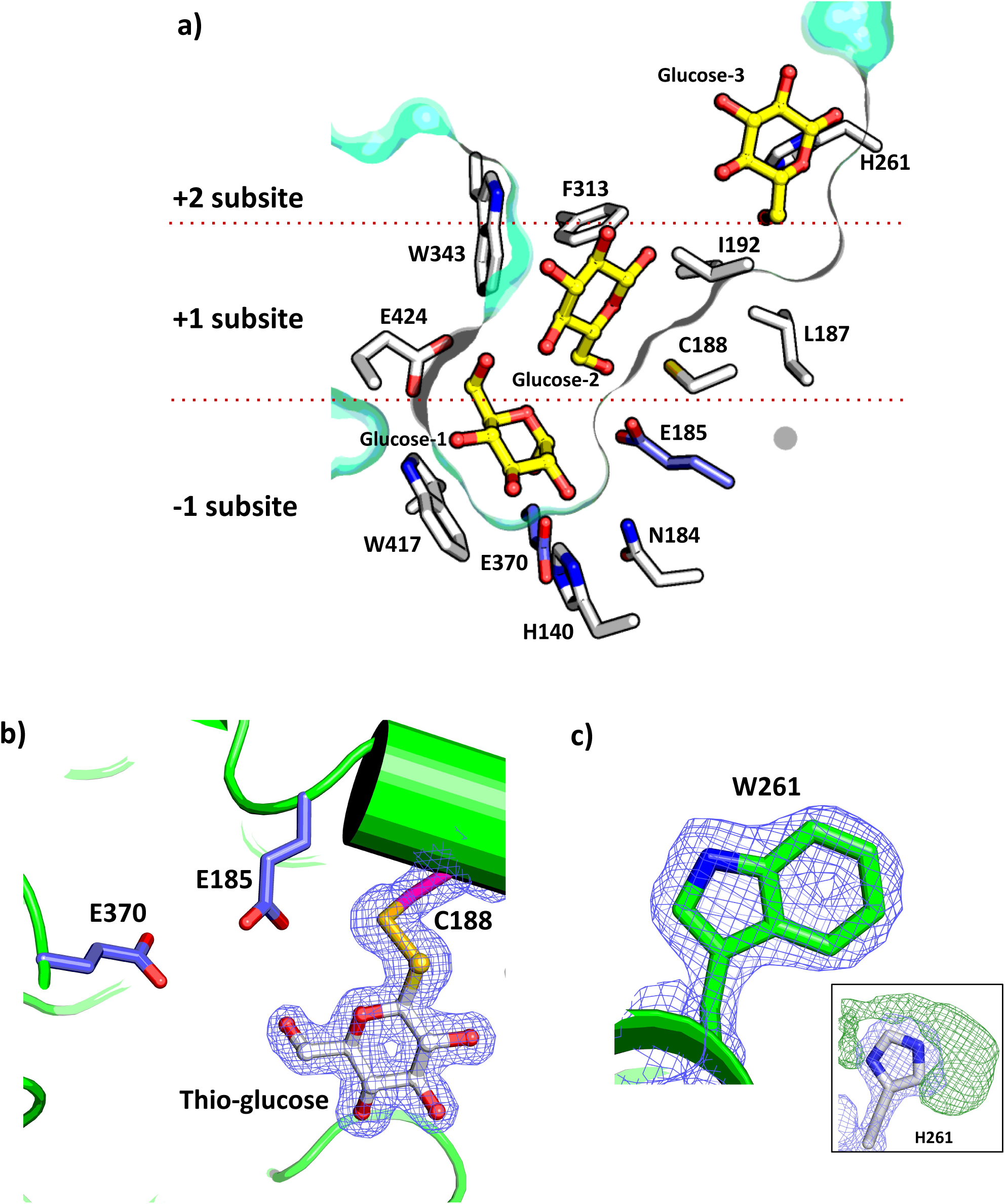
Understanding the molecular details of sugar-binding in the catalytic crater of UnBGl1. **a)** The crater is shown as surface (colour: greencyan) and lining representation, glucose molecules are shown in ball and stick (yellow carbon). Two catalytic glutamates are shown in purple sticks and all other residues are shown in white sticks. The −1, +1, +2 subsites are divided by dotted red lines. **b)** The 2*F_o_ - F_c_* electron density map (slate mesh) contoured at 1σ level around the thio-glycosidic enzyme intermediate is shown. Catalytic glutamates are shown in purple stick and cysteine residue is shown as magenta stick whereas thio-glucose is shown in ball and stick representation. **c)** The 2*F_o_ - F_c_* electron density map (blue mesh) contoured at 1 σ level indicates presence of tryptophan at 261^st^ position. Inset represents the omit *F_o_ - F_c_* electron density map (green mesh) suggesting the presence of tryptophan residue at 261^st^ position.

The third glucose molecule observed at the crater’s entry site was identified to present as the same site of glucose captured in cellobiose complexed structure (Supplementary Figure S3c), suggesting that this is a conserved glucose binding site. This third glucose binding site is assigned as the +2 subsite, which has close contact with the histidine residue at the 261^st^ (H261) position. Figure 4a shows all residues in the crater at different glucose binding subsites. In order to improve the glucose tolerance level of a GH1 β-glucosidase, it is essential to explore the product binding sites for modifying them to reduce the glucose accumulation in the crater at high glucose concentrations in the bulk solvent. Increasing the hydrophobicity of the surface of the crater can obstruct the entry as well as binding of glucose to the active site^5,13^. It’s a well-known fact that the aromatic residues play important role in the recognition of sugar substrates and may facilitate to increase the specificity of the enzyme^30,31^. Also, the aromatic residues at the entrance of the crater are responsible for substrate binding^32^. With this structural rational, a variant of UnBGl1 was created by replacing H261 with tryptophan and the crystal structure of UnBGl1_H261W variant in apo form was determined at 1.8 Å resolution (PDB ID. 9J43) (Table 1: part II). The electron density confirms the mutation as the side chain of W261 is well defined in the map (Figure 4c). Further the kinetic and biochemical characterizations, as well as structural studies of these variants as glucose bound complex were performed.

### Glucose tolerance and kinetic properties of UnBGl1 variants

Several variants of UnBGl1, were created targeting the residues in +1 and +2 subsites in order to improve the glucose tolerance of this enzyme. Biochemical properties of the variants were determined using the pNPG hydrolysis following the same protocol as reported for the native UnBGl1. UnBGl1_C188V did not change biochemical properties and had an optimal pH of 6.0 with an optimum temperature of 55 °C (Supplementary Figure S4c, S4d). However, UnBGl1_C188V showed difference in its kinetic properties (Table 2) (Supplementary Figure S4e) and has reduced affinity towards pNPG with *K*_m_ = 7.35 ± 3.0 mM. Maximum hydrolysis of cellobiose was seen at pH 6.5, and almost the same amount of activity was observed at pH 6.0 (Supplementary Figure S4f). Kinetic properties for cellobiose hydrolysis were determined at its optimum temperature 45 °C (Supplementary Figure S4g). As saturation of the UnBGl1_C188V mutant was not achieved (Supplementary Figure S4h), the kinetic properties of the enzymes for cellobiose hydrolysis could not be determined.

**Table 2:** Kinetic parameters of the native and variants of UnBGl1 towards hydrolysis of pNPG and cellobiose.

| Enzyme | $V_{\max}$<br>($\mu\text{mol min}^{-1}\text{mg}^{-1}$) | | $K_m$<br>(mM) | | $k_{\text{cat}}$<br>($\text{s}^{-1}$) | | $k_{\text{cat}}/K_m$<br>($\text{mM}^{-1}\text{s}^{-1}$) | | Glucose<br>Tolerance<br>(M) |
| --- | --- | --- | --- | --- | --- | --- | --- | --- | --- |
|  | <u>pNPG</u> | <u>Cellobiose</u> | <u>pNPG</u> | <u>Cellobiose</u> | <u>pNPG</u> | <u>Cellobiose</u> | <u>pNPG</u> | <u>Cellobiose</u> |  |
| *Native UnBGl1* | $36.14 \pm 0.50$ | $19.74 \pm 0.37$ | $0.92 \pm 0.05$ | $34.61 \pm 2.4$ | $31.62 \pm 0.57$ | $17.27 \pm 0.42$ | $34.36 \pm 11$ | $0.47 \pm 0.17$ | 1.0 |
| *UnBGl1_C188V* | $17.23 \pm 1.5$ | ND | $7.35 \pm 3.0$ | ND | $15.07 \pm 1.31$ | ND | $2.05 \pm 0.43$ | ND | 2.5 |
| *UnBGl1_H261W* | $89.18 \pm 1.36$ | $62.98 \pm 1.1$ | $0.65 \pm 0.04$ | $22.87 \pm 1.1$ | $78.03 \pm 1.19$ | $55.10 \pm 0.96$ | $120.05 \pm 29$ | $2.40 \pm 0.87$ | 1.6 |
| *UnBGl1_C188V_*<br>*H261W* | ND | ND | ND | ND | ND | ND | ND | ND | 2.5 |

ND = Not Determined

As mentioned earlier, we created another variant UnBGl1_H261W, by introducing mutation at +2 subsite, where glucose complexed structure showed alternate conformation of W261 (Supplementary Figure S5a and S5b). UnBGl1_H261W showed activity in a wide range of pH from pH 5.0 to 7.0 and retained almost 80 % of its ability to hydrolyse pNPG, considering the 100 % activity at pH 6.5 (Supplementary Figure S5c). UnBGl1_H261W showed maximum hydrolysis of pNPG at 55 °C and almost the same activity at 50 °C (Supplementary Figure S5d) with *K*_m_ of 0.65 ± 0.04 mM (Supplementary Figure S5e) (Table 2). Similarly, for cellobiose hydrolysis this variant enzyme showed activity within a short range of pH from pH 5.5 to 6.5. Activity at pH 5.5 was maximum and considered as 100 %, whereas pH 6.0 and 6.5 the enzyme has 90 % activity (Supplementary Figure S5f). This variant has same temperature optimum as of native UnBGl1 (Supplementary Figure S5g). UnBGl1_H261W has improved kinetic properties than native and UnBGl1_C188V. UnBGl1_H261W has shown increased affinity towards cellobiose with *K*_m_ of 22.87 ± 1.1 mM (Supplementary Figure S5h). The kinetic parameters of all variants of UnBGl1 are mentioned in the Table 2.

UnBGl1_C188V showed increased activity with a glucose tolerance level of about 2.5 M (Figure 5a). However, the kinetic properties were not improved (Table 2). Meanwhile, the UnBGl1_H261W variant exhibited improved kinetic properties for hydrolysing pNPG and Cellobiose (Table 2). The glucose tolerance of this variant is increased to 1.6 M (Figure 5b). As +1 subsite modification showed improved glucose tolerance and +2 modification showed improvement in kinetic properties therefore, a new variant of UnBGl1, referred as UnBGl1_C188V_H261W, was created by combining the modifications of these subsites (Supplementary Figure S6a). This double mutant gets stimulated by less concentration of glucose. Figure 5c illustrates an increase in enzyme activity with an increase in glucose concentration. UnBGl1_C188V_H261W has 2.5-fold higher activity at 250 mM of glucose, and at further glucose concentration, the enzyme started to show reduced activity (Figure 5c). Notably, compared to all variants of UnBGl1, the UnBGl1_C188V_H261W has the highest glucose tolerance of 2.5 M (Figure 5d). Meanwhile, its biochemical properties remain the same as those observed for the native UnBGl1 using pNPG hydrolysis (Supplementary Figure S6b, S6c). However, the kinetic parameters for hydrolysing pNPG could not be determined as pNPG is not soluble at higher concentrations (Supplementary Figure S6d). For cellobiose hydrolysis, pH optimum has shifted to 6.5, retaining more than 80% of activity at pH 6.0 (Supplementary Figure S6e) and reduced the optimum temperature for hydrolysis to 45 °C (Supplementary Figure S6f) the kinetic properties for cellobiose hydrolysis were also not determined as cellobiose is not soluble at higher concentration (Supplementary Figure S6g).

**Figure 5:**
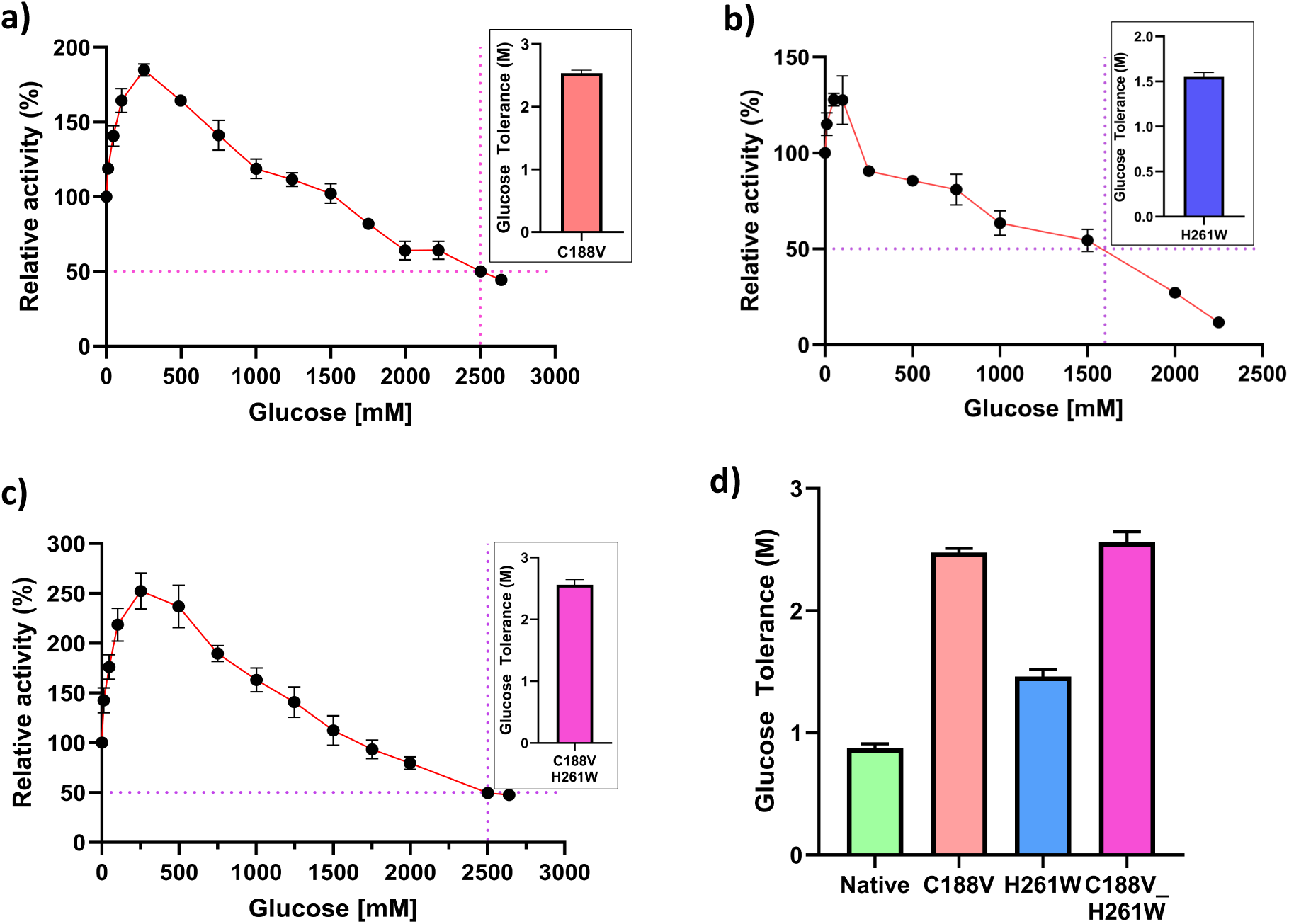
Glucose tolerance level of all the UnBGl1 Variants. Data points of the relative activity are shown in black solid circle and connected with red line. The concentration of the glucose that causes 50 % loss of the relative activity, is considered as glucose tolerance (indicated by purple dotted lines). **a)** The graph depicts glucose tolerance level of UnBGl1_C188V. Inset is another representation of glucose tolerance through bar graph. **b)** The graph depicts glucose tolerance level of UnBGl1_H261W. Inset is another representation of glucose tolerance through bar graph. **c)** The graph represents the glucose tolerance level of UnBGl1_C188V_H261W. Inset is a bar representation of glucose tolerance indicating highest glucose tolerance. **d)** Bar graph of comparison of glucose tolerance level of native UnBGl1 with variants. All experiments were performed in triplicates (n = 3) with error bar representing ± SEM.

### Evaluation of UnBGl1 variants for their industrial use

Simultaneous saccharification and fermentation (SSF) is widely used method for bioethanol production. The enzymatic hydrolysis and microbial fermentation occur in a single process and so saves the cost of extra vessel. Preparation of soluble fermentable sugars and fermentation of these sugar generally happens at a temperature of 25 °C - 30 °C for 6 - 72 hrs.^33^, therefore it is essential to have active and stable β-glucosidase for bioethanol production.

Further enzyme stability from sugarcane bagasse was monitored in the crude extract (cell lysate of expressed UnBGl1 variants cells). All the variants are stable at room temperature in pH 6.0 for 7 days (Figure 6a). On the 7^th^ day native UnBGl1, UnBGl1_C188V, UnBGl1_H261W, UnBGl1_C188V_H261W retained 61%, 65%, 84%, 68% of activity respectively (Figure 6a). Although all the variants have different activity towards cellobiose, they are stable for 7 days. Also, all the UnBGl1 variants were active when blended with a cellulase cocktail from *T. reesei* (Celluclast). Native UnBGl1 and UnBGl1_H261W, when blended with Celluclast, showed higher amount of glucose release of around 48 mM and UnBGl1_C188V, UnBGl1_C188V_H261W released 45 mM of glucose after 55 hrs. by hydrolysing sugarcane bagasse. Whereas, Celluclast only showed release of glucose of around 30 mM (Figure 6b). Further structural studies of UnBGl1 variants were done to understand the mechanism behind the improvement in glucose tolerance.

**Figure 6:**
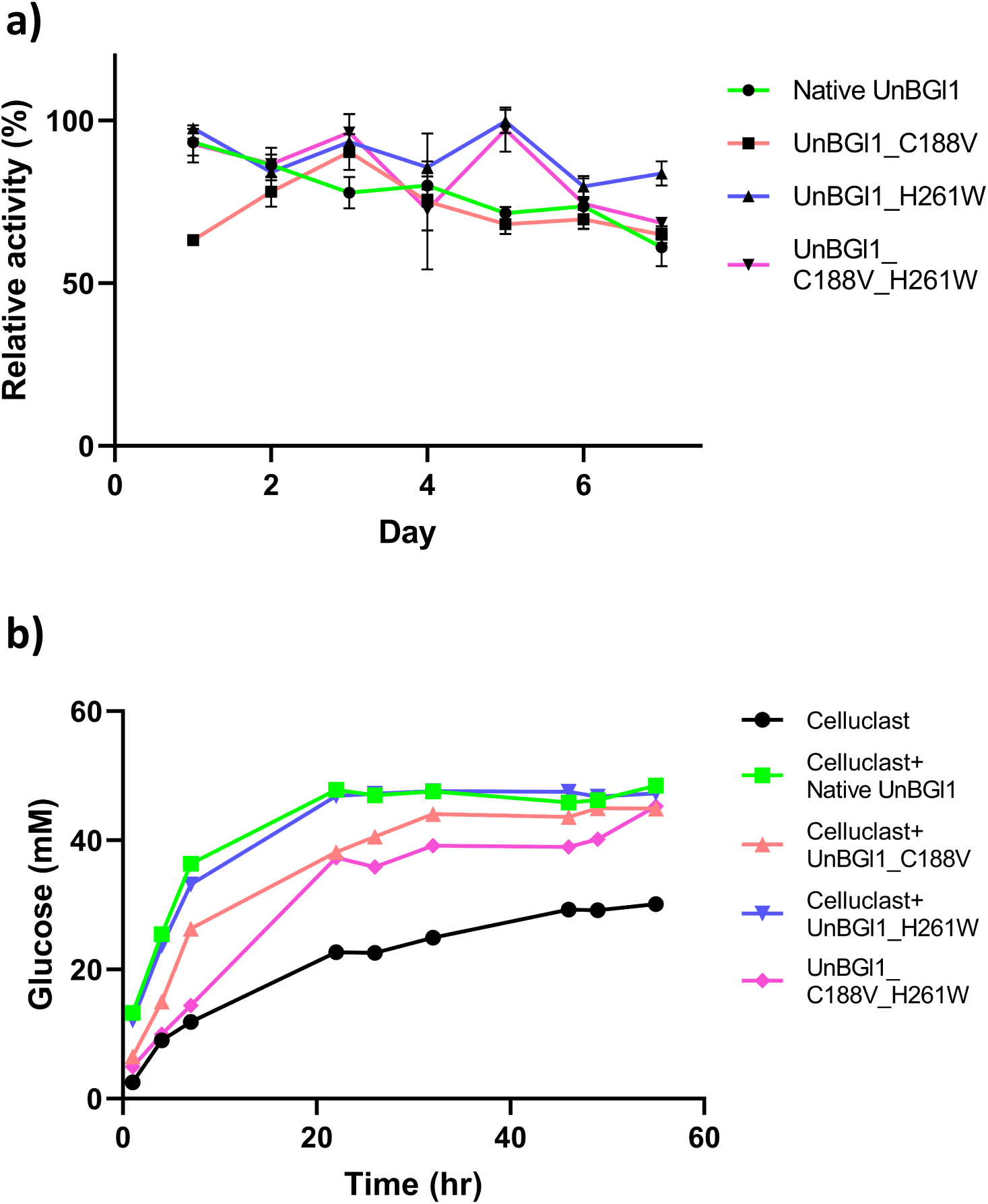
Evaluation of the properties of UnBGl1 variants as per the industrial requirements. **a)** Assessment of the stability and activity of UnBGl1 variants over the period of 7 days in cell lysate. **b)** Glucose release profiles of the UnBGl1 variants after blending the enzymes with industrially used β-glucosidase Celluclast.

### Structural relevance with the glucose tolerance

We determined multiple high-resolution crystal structures (Table 1: Part I and Part II) of the UnBGl1 variants as apo and complexed with glucose (Figure 7) to understand the molecular basis of improved glucose tolerance of these variants. We also solved the cellobiose bound structure of UnBGl1_H261W. The glucose complexed native UnBGl1 and UnBGl1_C188V structures do not show any overall structural difference, and their superposition produces a r.m.s.d value of 0.108 Å. The UnBGl1_C188V structure has only two glucose molecules in its crater, one at the −1 subsite and another at the +2 subsite (PDB ID. 9J42) (Figure 7a). As expected, glucose at the +1 subsite is absent in the UnBGl1_C188V structure (Figure 7a). This might be primarily because of loss of the interactions between glucose at the +1 subsite and the cysteine due to its replacement by a valine. Loss of interactions of glucose in the +1 subsite aided in reducing the glucose binding ability of the crater leading in reduction of glucose accumulation, leading to the increase of the glucose tolerance level (2.5 M) (Figure 5a) of UnBGl1_C188V as compared to the native UnBGl1 (0.9 M) (Figure 1f).

**Figure 7:**
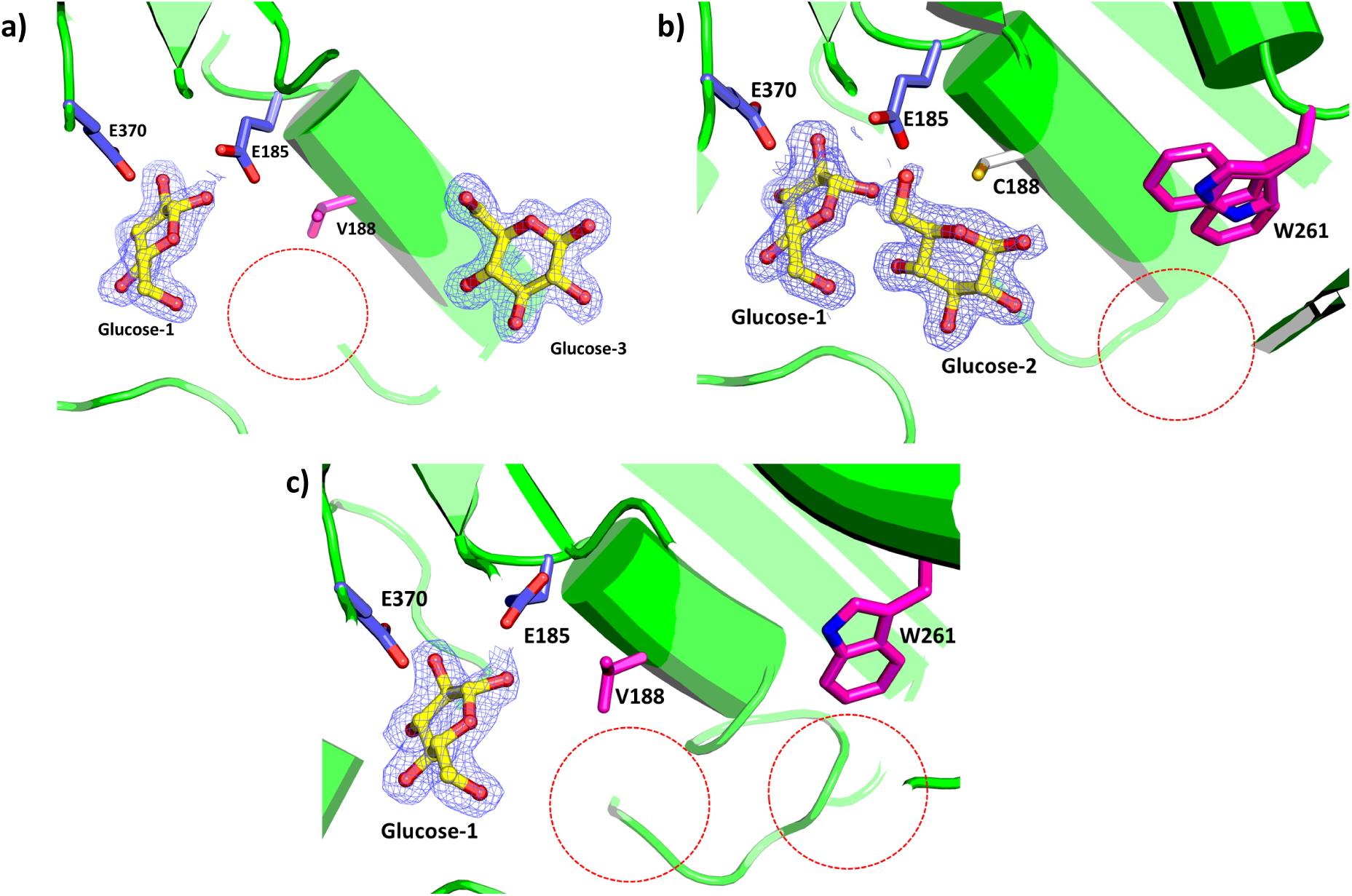
Molecular basis of improvement of the glucose tolerance of UnBGl1 variants. Zoomed in view of the active sites of the UnBGl1 variants showing the glucose binding. Catalytic glutamates are shown in purple stick, mutated residues (V188, W261) are shown in magenta stick, glucose molecules are shown as ball (red) and stick (yellow) representation. Absence of glucose molecules in the active site crater of these mutants as compared to the native enzyme (Figure 2e) is marked with dotted red circle. **a)** 2*F_o_ - F_c_* map contoured at 1 σ level illustrating the electron density (blue mesh) around two bound glucose molecules in UnBGl1_C188V variant structure. The glucose molecule near V188 is absent in this mutant. **b)** 2*F_o_ - F_c_* map contoured at 1.0 σ illustrating the electron density (blue mesh) around two bound glucose molecules in UnBGl1_H261W variant structure. The glucose molecule near W261 is absent, no electron density for glucose was observed near W261. **c)** The 2*F_o_ - F_c_* map contoured at 1 σ level showing an electron density (blue mesh) around a single glucose molecule near active site and absence of electron density of 2^nd^ and 3^rd^ glucose molecule.

The glucose tolerance of the UnBGl1_H261W was observed to be 1.6 M (Figure 5b) which is not greater than that of UnBGl1_C188V but much higher than the native enzyme. However, the kinetic properties of UnBGl1_H261W have improved as compared to the native and C188V variant (Table 2). The high-resolution glucose complexed UnBGl1_H261W structure (PDB ID. 9J44) provided insights into the active site crater to understand the basis for increased glucose tolerance level of this variant. The structure showed absence of a glucose molecule at the +2 subsite (Figure 7b). Further electron density analysis revealed the double confirmation of W261 (Supplementary Figure S5a, S5b). Flexible nature of the bulky hydrophobic side chain of W261 might be one of the reasons for the absence of a glucose at the +2 subsite. Decrease of glucose affinity due to the presence of a hydrophobic residue at the entry site (+2 subsite) of UnBGl1_H261W crater contributed to increase glucose tolerance level of this enzyme.

Structure of glucose complexed UnBGl1_C188V_H261W double mutant was refined at 2.0 Å (PDB ID. 9J4A) (Table 1: part II). Double mutant showed the absence of glucose molecules at +1 and +2 subsites and a single molecule of glucose was present only at −1 subsite (Figure 7c). This suggest that it requires high concentration of glucose to get trapped into the crater of double mutant UnBGl1. Our structural and biochemical data clearly indicate that incorporation of hydrophobic residues at +1 and +2 subsites of UnBGl1 has aided to improve glucose tolerance of this enzyme.

## Discussion

GH1 β-glucosidases play pivotal role in process of degradation of cellulose in smaller fermentable glucose units by hydrolysing β,1-4 linkage^18^. These enzymes possess feedback inhibition property, emphasizing the necessity for β-glucosidases with enhanced glucose tolerance to ensure efficient and continuous cellulose breakdown^34^. In this investigation, a β-glucosidase from soil metagenome (UnBGl1) has been studied extensively. We have determined 12 high-resolution crystal structures (including apo, substrate-bound, product-bound, and inhibitor-bound forms), of UnBGl1 and its variants. Taken together our findings, the strategies and important structural features to be considered for the rational design to improve glucose tolerance of GH1 family β-glucosidases are discussed.

### Understanding cellobiose hydrolysis mechanism through crystallographic snapshots

For native β-glucosidases, no structure is reported so far that shows substrate binding and presence of a covalently linked glycosyl intermediate in the active site. Here, we present the first-ever crystal structure of β-glucosidase in its native state complexed with the substrate, cellobiose, precisely at the pre-hydrolytic state (Figure 2c). In case of UnBGl1, E185 and E370 are catalytic residues that carry overall hydrolytic reaction. The binding and hydrolysis of cellobiose have been studied through computational techniques. The mode of cellobiose binding seen in UnBGl1 catalytic pocket is similar to that observed previously by the quantum mechanics/molecular mechanics (QM/MM) studies on hydrolysis of laminaribiose (β-1,3 linked) and cellobiose by a plant β-glucosidase (BGul1)^35,36^. During the initial binding phase, the enzyme’s active site adjusts to the binding of cellobiose, making it easy for the substrate to be catalytically processed. Substrate binding further induces conformational changes in UnBGl1, aligning cellobiose appropriately within the catalytic site. QM/MM calculations have demonstrated how the enzyme dynamically adapts the positioning of substrates to optimize conditions for nucleophilic attack, thus underlining structural adaptability during catalysis. The same orientation of cellobiose is seen in the UnBGl1 active site. This state is stabilized by various hydrogen bond interactions between cellobiose and active site residues. Hydrogen bonding interactions (Supplementary movie S3) seen in the crystal structure of cellobiose bound UnBGl1 are comparable to those reported for BGul1. The correct orientation of cellobiose is necessary to achieve transition states and further formation of glycosyl intermediate. This study is the first report of capturing a glycosyl intermediate in native and active β-glucosidase. So far, no experimental structural evidence of pre-hydrolytic state of substrate and glycosyl intermediate has been reported for β-glucosidases. The structural snapshot of cellobiose complexed UnBGl1 elucidates critical substrate-enzyme interactions and sheds light on the initial steps of the enzymatic reaction. Also, the structure of a reaction intermediate where the substrate forms a covalent bond with the nucleophilic glutamate residue in the active site, offers a deeper understanding of the transition-state stabilization during catalysis.

### Glucose binding subsite mutations and corresponding modulation in kinetic properties in UnBGl1

Insufficient β-glucosidase activity is the reason for the accumulation of cellobiose in the enzymatic cellulose hydrolysis process, which results in the inhibition of endo- and exo-glucanases^16^. Therefore, identifying the critical residues that can enhance the efficiency of β-glucosidases activity is highly needed. Determination of the kinetic parameters of β-glucosidases is essential for evaluating their efficiency and potential industrial applications. Here, we investigated the kinetic properties of all the variants of UnBGl1 against their enzyme property to hydrolyse pNPG and cellobiose (Table 2). Most of the β-glucosidase enzymes have an optimum temperature above 40 °C and exhibit the maximum activity in the pH range of 4.0 - 6.5; as documented by Teugjas et al.^37^. UnBGl1 and all of its variants have the same property with an optimum temperature range of 45 - 55 °C and pH range of 4.0 - 6.5 for both pNPG and cellobiose hydrolysis. Efficient industrial cellulose conversion to simple sugar like glucose requires highly stable and active enzymes. A process like simultaneous saccharification and fermentation (SSF) uses active and stable enzymes at lower temperatures^38^ of around 30 °C - 40 °C^39^. UnBGl1 satisfies the requirements of SSF process with enzyme stability of 48 hrs at 30 °C in pH 5.5 and 6.0 (Figure 1d). All the UnBGl1 variants were stable in the cell lysate for 7 days at room temperature (Figure 6a), and therefore the use of these variants in the industry will save a lot of protein purification and storage expenses. When kinetic parameters were compared of native enzyme with +2 subsite H261W variant, native UnBGl1 showed more affinity and catalytic efficiency towards both pNPG and cellobiose. Meanwhile, the +1 subsite C188V variant drastically changed the kinetic behaviour of the enzyme and could not achieve complete substrate saturation. A similar observation has already been reported where a β-glucosidase from *Themococcus sp.* (O08324) was modified by introducing various mutations at +1 and +2 subsite^40^. H0HC94, a β-glucosidase from *Agrobacterium tumefaciens* 5A showed lower affinity after L178E mutation was introduced at +1 subsite towards both pNPG and cellobiose^41^. Based on our structural analysis, we believe that +1 subsite plays an important role in maintaining the kinetic properties of GH1 β-glucosidases. Extreme care and through structure as well as sequence analysis have to be done if there is a need to modify the +1 subsite. It is important to determine the enzyme’s activity when it is used in cellulase cocktails. For that we used Celluclast (an enzyme cocktail from *T. reseei* that does not have β-glucosidase) blended with variants of UnBGl1 and used to hydrolyse bagasse. UnBGl1 variants can release more glucose from bagasse when added to the cellulase cocktail (Figure 6b). Therefore, our data suggest that these variants of UnBGl1 are suitable candidates for the SSF process for ethanol production.

### Structural basis of the glucose stimulation and tolerance in GH1 β-glucosidases

Based on the effect of glucose concentration, β-glucosidases are classified into four groups that: (i) get inhibited at lower concentrations of glucose, (ii) are glucose tolerant, (iii) get stimulated at low concentrations of glucose, (iv) do not undergo inhibition at high concentration of glucose^42,43^. In this study, we have selected UnBGl1, a β-glucosidase that shows glucose stimulation at lower glucose concentrations (Figure 1f, 5a, 5b, 5c). Our biochemical studies indicate that the glucose can bind to the stimulatory sites of the enzyme active site crater up to a specific concentration, which is 100 mM for the native and UnBGl1_H261W, whereas in the case of UnBGl1_C188V and UnBGl1_C188V_H261W, it has increased to 250 mM. However, beyond that stimulatory glucose concentration, all variants start to show inhibition. Compared to the native enzyme, all the variants have showed improved glucose tolerance. Although, UnBGl1_H261W with +2 subsite mutation did not show any effects on the glucose stimulation activity, both single and double mutant variants having cysteine to valine substitution at +1 subsite exhibited an increase in the stimulatory glucose concentration and the stimulatory effect. It is evident that the +1 subsite plays a vital role in the glucose stimulation, but the mechanism of such effect is still unclear. It has been postulated that this effect may arise from the geometric configuration of specific binding sites, where bound glucose enhances substrate cleavage activity through transglycosylation or other mechanisms, such as allosteric modulation^44^. Our extensive structural data and analysis of multiple ligand complexed UnBGl1 structures, two predominant allosteric glucose binding sites (+1 and +2 subsites) away from the catalytic centre have been identified. However, modifying those two allosteric sites (in UnBGl1_C188V and UnBGl1_H261W) did not diminish the glucose stimulatory effect. Therefore, the allosteric binding of glucose may not be the reason for stimulatory effect of β-glucosidases.

Glucose influences the active site environment by altering the water matrix and steric geometry in the substrate channel or at the catalytic centre, subsequently affecting substrate binding. Additionally, glucose binding outside the active site, potentially in the middle of the substrate channel or at its entrance, can increase glucose tolerance^44^. However, our structural and biochemical data indicate that loss of interaction of glucose from +1 and +2 subsites increases glucose tolerance in UnBGl1 variants. All three variants, UnBGl1_C188V, UnBGl1_H261W and UnBGl1_C188V_H261W, that showed loss of interaction of glucose from the middle and entry site of the crater exhibited significant improvement in glucose tolerance to 2.5 M, 1.6 M and 2.5 M, respectively. These values are higher than the glucose tolerance reported for many β-glucosidases^37^ so far. The high-resolution crystal structures clearly indicate that glucose binding sites are very specific in the case of UnBGl1. That proves, glucose binding in the UnBGl1 crater is not non-specific. Glucose binding to +1 and +2 subsites are responsible for low glucose tolerance of the enzyme due to blockage of the entry channel resulting in non-accessibility of the substrate to catalytic glutamates (E185, E370) at the −1 subsite for hydrolysis. Removal of the interactions of glucose from the +1 and +2 subsites in the crater is essential to facilitate the entry of the substrate cellobiose to the catalytic centre. Modifications at the +1 subsite by introducing valine at the 188^th^ position relieved glucose from +1 subsite (Figure 8b), leading to increased glucose tolerance. Introduction of a hydrophobic residue at +2 subsite has also shown loss of interactions with glucose at +2 subsite (Figure 8c) leading to increase in glucose tolerance. Creating a double mutant by combining +1 and +2 subsite modifications has led to further increase in glucose tolerance (Figure 8d).

**Figure 8:**
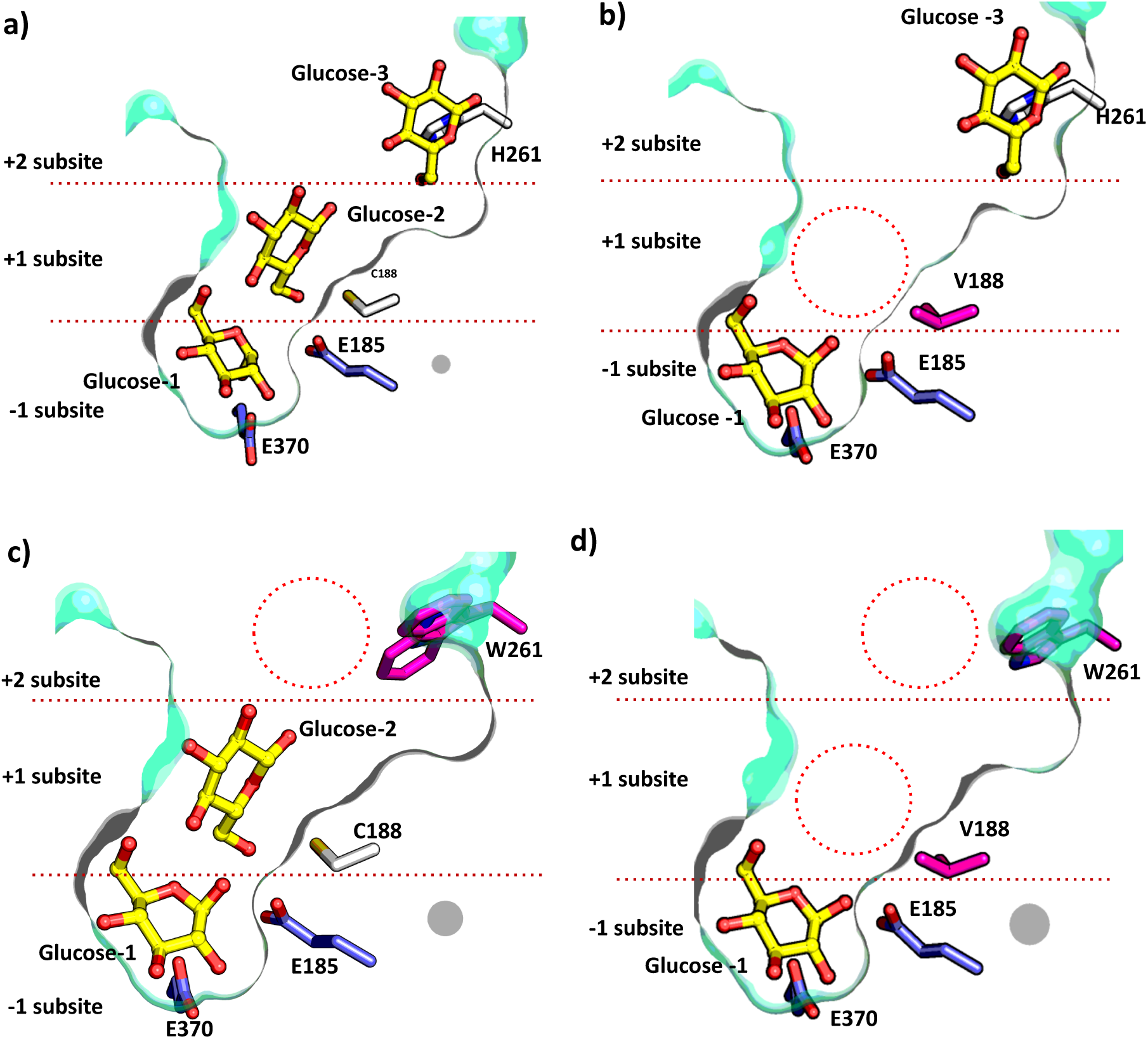
Effect of mutations on the conserved glucose binding sites in UnBGl1 variants. Crater of all the variants is shown in surface representation, catalytic glutamates are in purple stick and other native residues are in white sticks; whereas mutated residues are shown in magenta sticks. All glucose molecules are illustrated as ball and stick. Absence of glucose molecules in the active site crater of these mutants as compared to the native enzyme is marked with dotted red circle. **a)** Three glucose molecules bound to the subsites of native UnBGl1 crater. **b)** Absence of second glucose from middle of the crater in UnBGl_C188V due to C188V mutation at +1 site. **c)** Absence of third glucose from entry site of the crater in UnBGl1_H261W due to H261W mutation at +2 site. **d)** Absence of two glucose molecules at +1 and +2 subsites in UnBGl1_C188V_H261W.

### Role of residues in catalytic crater towards increasing glucose tolerance of GH1 β-glucosidases

As GH1 β-glucosidases share a highly similar structural fold, the findings from our structural studies on UnBGl1 can be applicable to other enzymes within the same family. Through structure-based sequence alignment with glucose-tolerant β-glucosidases from the Betagdb database^42^ and few other β-glucosidases, we have identified residues that could be modified to boost glucose tolerance, as well as those that should remain untouched in GH1 β-glucosidases. The supplementary movie S4 illustrates the residues involved in formation of the catalytic core of UnBGl1, which are highly conserved across most β-glucosidases. We believe altering these amino acids could negatively impact enzyme activity due to their proximity to the catalytic residues. Characteristic NEP and ENG motifs, which are essential features of GH1 β-glucosidases, are also part of this core. R96 interacts with E370 (salt bridge) and N184 (hydrogen bond), which are part of the catalytic core and contribute to the stability of the active site (Supplementary Figure S7a) in UnBGl1. Whereas Q39 forms hydrogen bond with glucose or cellobiose, as observed in crystal structure (Supplementary Figure S7b, S7c). These hydrogen bonds are essential for suitable conformation of cellobiose in the active site for hydrolysis. H140 is located at the crater’s base and closes the catalytic crater as shown in the supplementary movie S4. Additionally, W141, W417 and W425 surround the catalytic core of the enzyme and maintain the hydrophobic environment. Y312 and E424 residues from +1 subsite are present at hydrogen bonding distance from glucose and cellobiose present at −1 subsite (Supplementary Figure S7d, S7e). The residues Q39, R96, H140, W141, N184, E185, P186, E370, W417 and W425 are present at −1 subsite, Y312 and E424 present at +1 subsite are conserved across all glucose tolerant β-glucosidases (Supplementary Figure S8a). Modifying these residues could adversely affect cellobiose binding and enzyme activity, making such alterations unwise for improving glucose tolerance in GH1 β-glucosidases.

Our structural and biochemical studies have underscored the critical role of C188 and H261 residues in improving glucose tolerance. Multiple sequence alignment of glucose tolerant GH1 β-glucosidases showed the presence of aromatic residues at 187^th^ site in +1 subsite. Most of the β-glucosidases have tryptophan residue at this site so introducing tryptophan at 187^th^ position can increase the glucose tolerance. This was demonstrated by mutating G168 to tryptophan in β-glucosidase from *Acetivibrio thermocellus* (AtGH1), which increased in glucose tolerance to 820 mM^45^. G168W variant of AtGH1strucutrally aligns with L187, therefore L187 to tryptophan mutation can increase glucose tolerance of UnBGl1. Similarly, at 192^nd^ and 199^th^ position Ile and Met are present in UnBGl1 whereas most glucose tolerant β-glucosidases have Leu and His at this position (Supplementary Figure S8b). Generating I192L and M199H variants of UnBGl1 would increase the glucose tolerance. Based on our structural and bioinformatic studies, it is evident that L187W, C188V, I192L, M199H, and H261W modifications at the +1 and +2 subsites can enhance glucose tolerance. This study lays a robust foundation for the rational design of GH1 family β-glucosidases, enabling the development of enzyme variants with improved performance under industrially relevant conditions. Such advancements are poised to contribute substantially to the economic viability and efficiency of bioethanol production processes.

## Materials and Methods

### Cloning, mutagenesis, protein expression and purification

The codon optimized sequence of *UnBGl1* gene was synthesized from GeneArt, Invitrogen (USA) to express in *Escherichia coli* (*E. coli*) strain BL21(DE3). *UnBGl1* gene was amplified by PCR using gene-specific primers with help of Q5 polymerase enzyme (NEB) and was cloned into *NdeI* and *XhoI* sites by using T4 DNA ligase enzyme (ThermoFisher Scientific) of pET43.1b to obtain C-terminal His_6_ tag protein. The recombinant plasmid was transformed in *E. coli* BL21(DE3) for over-expression of UnBGl1. These cells were grown at 37 °C for 16 hrs. In 50 ml Luria-Bertani (LB) media containing 100 mg/ml ampicillin. 20 ml of this grown culture was used to inoculate in 1.5 lit LB media with the same ampicillin concentration. *E. coli* BL21 (DE3) cells were grown in 3 lit LB medium to obtain a high amount of purified protein. Cells were grown at 37 °C till OD at 600 nm reaches 0.6 - 0.8. The cells were induced with 0.4 mM IPTG for overexpression of the protein, and cells were grown at 24 °C for 16 hrs. The media was centrifuged at 8000 rpm for 10 minutes to collect the cell pellet. After harvesting the cells, 5x volume of sodium phosphate buffer (50 mM Na-PO_4_, 400 mM NaCl, and pH 7.4) was added to the pellet to dissolve. Cell disruption was performed with ultrasonication at 4 °C and centrifuged at 16000g for 30 minutes. A 0.4 *μ*m PVDF membrane was used to filter the supernatant.

The supernatant was loaded onto a 5ml His Trap FF Ni-NTA affinity column equilibrated with sodium phosphate buffer using FPLC (ÄKTAprime Plus, GE Healthcare) at a flowrate 1 ml/minute. UnBGl1 was eluted using equilibration buffer containing increasing concentrations of imidazole (25 mM, 75 mM, 125 mM, 187 mM and 100 mM). The maximum amount of protein was eluted with 75 mM imidazole. Eluted protein was further pooled and concentrated to 2 ml using an Amicon ultrafiltration concentrator of 10 kDa cutoff. The concentrated protein was purified by a pre-equilibrated (50 mM Na-PO_4_, 50 mM NaCl, pH 7.4) Superdex 200 16/600 (GE Healthcare) gel filtration column. Peak fractions were collected, and the purity of the enzyme was analysed by running SDS-PAGE.

Further for preparation of site directed mutants (SDMs), site specific primers (Supplementary Table S1) were designed. Recombinant native UnBGl1 plasmid was amplified using site specific primers by Q5 polymerase enzyme for introducing C188V and H261W mutation. To create UnBGl1_C188V_H261W variant, UnBGl1_C188V plasmid was chosen as template and UnBGl1_H261W SDM primers were used for amplification by Q5 polymerase. The expression and purification of all the variants of UnBGl1 was done by following the protocol used for native UnBGl1.

### Enzyme Assay

Understanding the ideal reaction conditions for carrying out substrate hydrolysis is crucial. pH and temperature are the two most crucial parameters to comprehend in enzymatic activity. The enzymatic characterization of UnBGl1 was done using two substrates, p-nitrophenyl-β-D-glucopyranoside (pNPG) (Merck) and cellobiose (Sigma-Aldrich). All assays were conducted under optimal conditions to ensure reliable measurement of enzymatic activity.

For pNPG hydrolysis assay, a 40 mM stock solution was prepared in 50 mM sodium acetate and 50 mM sodium phosphate buffer of optimum pH. Reaction mixtures containing 40 µl of appropriately diluted pNPG solution and 450 µl of buffer were pre-incubated at optimum temperature for 5 minutes. 10 µl of appropriately diluted UnBGl1 was added to the pre-incubated reaction mixture and reaction was carried out on dry bath for 10 minutes. The reaction was terminated by addition of 500 µl of 0.2 M sodium carbonate and release of product p-nitrophenol (pNP) was quantified spectrophotometrically by measuring absorbance at 405 nm. For cellobiose hydrolysis assay, 350 mM cellobiose stock was prepared in 50 mM sodium acetate and 50 mM sodium phosphate buffer of optimum pH. Final reaction volume was set to 100 µl that contain 95 µl of appropriately diluted cellobiose and 5 µl of UnBGl1. The assay was carried out at optimum temperature for 30 minutes and terminated by heating at 90 °C for 10 minutes. After cooling, 2 µl of the reaction mixture was mixed with 200 µl Glucose-Oxidase Peroxidase (GOD-POD) reagent (Eco Pak Glucose 500, Accurex Biomedicals Pvt. Ltd.)^46^ in 96-well microtiter plate and incubated at 37 °C for 15 minutes. The developed pink colour, indicative of glucose production was measured spectrophotometrically at 505 nm using a microplate reader (Vantastar, BMG Labtech). Above mentioned protocol was used for characterization of all UnBGl1 variants.

Specific activities for both substrates were calculated as the amount of substrate hydrolysed (µmoles) per minute per milligram of enzyme and expressed in µmol/min/mg. Enzyme activities were measured in triplicates, and negative controls without enzyme were included to account for non-enzymatic hydrolysis. Standard calibration curves for pNP (405 nm) and glucose (505 nm) were used for accurate quantification of reaction products.

### Determination of optimum pH and optimum temperature

The reaction mixtures and substrates were prepared in 50 mM sodium acetate (pH 3.5, 4.0, 4.5, 5.0, 5.5) and sodium phosphate (pH 6.0, 6.5, 7.0, 7.5, 8.0) in these buffers These helped a comprehensive analysis of enzymatic performance under various pH conditions. For pNPG hydrolysis, 10 µl of purified UnBGl1 (1:200 dilution of 1 mg/ml) was used. Assay was carried at 55 °C for 10 minute and terminated by adding 0.2 M sodium carbonate. Maximum release of pNP was identified and considered as optimum pH for pNPG hydrolysis.

For cellobiose hydrolysis assay, 5 µl of 1 mg/ml UnBGl1 was used. Reaction was carried out in PCR machine at 50 °C for 30 minute and terminated at 90 °C. GOD-POD kit used to determine release of glucose, maximum glucose release was identified by spectrophotometric measurement at 505 nm. The value obtained at optimum pH was defined as 100 %.

Further for determining the optimum temperature for substrate hydrolysis, the reaction mixture and substrates were prepared in 50 mM sodium phosphate, pH 6.0 buffer. pNPG and cellobiose hydrolysis reactions were carried out in temperatures range of 25 °C - 80 °C.

For both assays, the activity at the optimal pH and temperature was normalized to 100% and served as the reference for relative activity comparisons across the tested pH and temperature range.

### Stability of native UnBGl1

The temperature stability of the native UnBGl1 was checked at 30 °C for 48 hrs. Cellobiose, the actual substrate of β-glucosidase, was used for performing stability tests. Experiments were performed in 50 mM sodium acetate (pH 5.5) and 50 mM sodium phosphate (pH 6.0) buffers. The reaction mixture was made such that it contained 990 µl of 350 mM cellobiose prepared in the buffer. 10 µl of native UnBGl1 (1mg/ml) was added to the reaction system. The cellobiose hydrolysis reaction was carried out at 50 °C for 30 minutes. The reaction was inhibited by placing reaction tubes in a boiling water bath for 15 minutes. 10 µl of an aliquot from the reaction mixture was mixed with 1 ml of GOD-POD reagent^46^. This assay system was incubated at 37 °C for 15 minutes. The pink colour was estimated spectrophotometrically at 505 nm for calculating enzyme activity.

Further, the cell lysate of all UnBGl1 variants were incubated at room temperature for 7 days at pH 6.0 and activity of the variants against cellobiose hydrolysis was measured each day as mentioned above.

### Measurement s of kinetic parameters

Enzyme kinetics studies of UnBGl1 variants were performed using its original substrate (cellobiose) and artificial substrate 4-nitrophenyl β-D-glucopyranoside (pNPG). To determine the kinetic parameters towards cellobiose hydrolysis, increasing concentration of cellobiose (3.5 mM to 332.5 mM) was incubated with the appropriately diluted enzyme at 50 °C for 30 min, and the reaction was inhibited by incubating at 90 °C for 10 minutes. Finally, to determine the amount of glucose released, 2 µl from the reaction mixture was taken, and 200 µl GOD-POD reagent was added and incubated at 37 °C for 15 min, and absorbance was measured at 505 nm. To determine the enzyme’s kinetic properties for hydrolysing the pNPG, increasing concentration of pNPG (50 to 40 mM) incubated with appropriately diluted enzyme at 55 °C for 10 min and the reaction was inhibited by adding 0.5 ml of 0.2M Na_2_CO_3_. The absorbance of the produced yellow colour was measured at 405 nm.

### Evaluation of glucose tolerance of UnBGl1 variants

The glucose tolerance level of the native and variants of UnBGl1 was determined by performing a pNPG hydrolysis assay in the presence of an increasing concentration of glucose (0 mM to 2.3 M). A fixed concentration of pNPG (40 µl of 40 mM) was incubated with fixed concentration of enzyme (10 µl of 1:200 diluted 1 mg/ml enzyme) and increasing glucose concentration. The reaction was carried at 55 °C for 10 minutes and inhibited by adding 0.2 M Na_2_CO_3,_ and absorbance was measured at 405 nm.

### Monitoring activity against sugarcane bagasse

The sugarcane bagasse was pretreated to separate the cellulose^47,48^. Obtained cellulose was further used for hydrolysis assay. An enzyme cocktail of cellulase from *Trichoderma reseesi* (Celluclast) that does not contain β-glucosidase was choose for this assay. The Celluclast was blended with the UnBGl1 variants in 1:1 ratio (Undiluted (neat) Celluclast: 2 mg/ml of UnBGl1 variant) to prepare new cocktail that can hydrolyse the cellulose to glucose. 50 µl of newly prepared enzyme cocktail was used to hydrolyse 20 mg/ml of bagasse prepared in sodium phosphate buffer (pH 6.0) and incubated at 30 °C for 55 hrs. Released glucose was measured by GOD-POD kit.

### Crystallization of UnBGl1 variants and crystal soaking experiments

Purified UnBGl1 was concentrated to 10 mg/ml in 50 mM sodium phosphate buffer containing 50 mM NaCl pH 7.4, using 10 kDa Amicon centrifugal unit. The concentrated protein was subjected to crystallization screening using Phoenix crystallization robot (Art Robbins Instruments) by sitting drop vapor diffusion method in 96-well Intelli-plates available at the “Protein Crystallography Facility”, Indian Institute of Technology, Bombay, India. For the crystallizations, different commercially available crystallization screens such as PEGS suite (Qiagen), PEGRx (Hampton Research), PEG/Ion (Hampton Research), Index (Hampton Research) and JCSG-plus (Molecular Dimensions) were used. Native UnBGl1 got crystallized after 90 days at 22 °C in a condition consisting of 0.056 M Sodium phosphate monobasic monohydrate, 1.344 M Potassium phosphate dibasic, pH 8.2 (condition 19 from Index - HR2- 144, Hampton research). Further optimization of crystallization was carried out by performing the hanging-drop vapor diffusion method using flat bottom 24 well polystyrene plates (Nest Biotech) and incubated at 22 °C. Different ratios of UnBGl1 to crystallization buffer (mother liquor), such as 1:1, 1:2, and 2:1 were tested to obtain high-quality crystals. The crystallization drops were set on silicon coverslips and placed over the reservoir having 500 µl of mother liquor. Once crystals were obtained, they were used for seeding experiments to enhance crystal growth. Seeding experiments successfully reduced the crystal growth time to 3 - 5 days and produced better quality crystals. The same protocol was applied for crystallizing all other variants of UnBGl1, ensuring consistency in crystal quality and reproducibility.

### Soaking experiments to obtain ligand complexed structures

As cellobiose hydrolysis is one step reaction and does not require any co-factor or activator for completion of reaction, co-crystallization is not a suitable method for obtaining substrate (cellobiose) complexed structure. Soaking experiment is often used to obtain protein-ligand complexes^49^, ligands can reach the binding site by diffusing through the solvent channels^50^. To confirm the catalytic activity of UnBGl1 in its crystalline form, initial soaking experiments were conducted using 40 mM p-nitrophenyl β-D-glucopyranoside (pNPG). A crystal of UnBGl1 (after washing multiple times with mother liquor) was transferred into the pNPG solution prepared in mother liquor, and the enzymatic activity was verified by observing yellow colour in the crystals and the surrounding solution.

Further soaking experiments were carried out with cellobiose at concentrations ranging from 50 mM to 350 mM. Native UnBGl1 and UnBGl1_H261W crystals were harvested using cryo-loops, gently washed with mother liquor, and soaked in cellobiose solutions for varying durations (10 seconds to 15 minutes) to determine optimal binding conditions. To capture a reaction intermediate, native UnBGl1 crystals were soaked in 300 mM cellobiose for 7 minutes. Additionally, native UnBGl1 crystals were soaked in 1 M glucose for 3 minutes to identify conserved glucose binding sites within the active site. And thio-cellobiose soaking was performed to find inhibitory sites of UnBGl1. Crystals were carefully handled to minimize physical damage during transfer and soaking procedures. Same protocol was followed to capture glucose bound structures of UnBGl1 variants. The soaking conditions are mentioned in Supplementary Table S2.

### X- ray diffraction data collection and processing

Diffraction data were collected by rotation method under liquid nitrogen cryo-conditions at 100 K, for that suitable cryoprotectants were prepared in mother liquor. Cryoprotectants used for freezing the crystals are mention in Supplementary Table S2. Crystals were carefully transferred into their respective cryoprotectants, mounted on a goniometer head, and flash-cooled under a liquid nitrogen stream at 100 K.

For glucose soaked native UnBGl1, UnBGl1_C188V, UnBGl1_H261W, cellobiose-soaked native UnBGl1, and thio-cellobiose-soaked native UnBGl1 crystals, diffraction datasets were collected using Cu*K*α X-ray radiation generated by a Rigaku Micromax 007HF generator equipped with an R-Axis IV++ detector at the Protein Crystallography Facility, IIT Bombay, India. Apo native UnBGl1, UnBGl1_C188V, UnBGl1_H261W and UnBGl1_C188V_H261W, glucose soaked UnBGl1_C188V_H261W and cellobiose soaked UnBGl1_H261W crystals were frozen in liquid nitrogen. The diffraction data of these crystals were collected using a synchrotron radiation X-ray source at beamline PX-BL21, RRCAT, Indore, India. The diffraction datasets of cellobiose soaked native UnBGl1 and glucose soaked UnBGl1_H261W crystals were processed using Xia2^51^ and Dials^52–54^ from CCP4Cloud Remote. Diffraction datasets of all other UnBGl1 variants were integrated using *XDS* and scaled using *XSCALE*^55^. The intensities were converted to structure factors with *F2MTZ* and *CAD* from *CCP*4^54^. Data collection statistics are represented in Table 1.

### Structure determination, model building and refinement

The native apo structure of UnBGl1 was determined using the molecular replacement (MR) method in *PHASER*^56^. The structure of a β-glucosidase from compost metagenome, Td2F2 (PDB ID. 3WH5)^57^ having 52.8 % amino acid sequence identity with UnBGl1, was used as a search model to compute initial phases. Matthews coefficient (2.23 Å^3^ Da^-1^)^58^ indicated one UnBGl1 molecule in asymmetric unit. After finding the correct solution, the template model was refined for 10 cycles using *REFMAC5*^59^. Initial model building was done in *COOT*^60^. Solvent molecules and ions were incorporated into the structure by identifying peaks in σ-A weighted *F_o_ - F_c_* electron density maps exceeding 3σ. These additions were carefully monitored to ensure a progressive reduction in *R_free_* and improvement of the overall stereochemistry. Final refinement was done by PDB-redo which automates model rebuilding and validation for optimal stereochemical and structural parameters^61^.

Structures of other variants of UnBGl1 were solved using apo native UnBGl1 (PDB ID. 9JLZ) as a template for molecular replacement. Ligand bound structures (including complexes with cellobiose, thio-glucose, glucose, and glycosyl intermediates) were resolved using rigid body refinement to derive initial phases. Following this, the protein components of the structures were subjected to initial refinement cycles to enhance the electron density maps of the ligands (cellobiose, thio-glucose, glucose, and glycosyl intermediates). Subsequently, the ligands were modelled into their corresponding σ-A weighted *F_o_ - F_c_* electron density maps, followed by additional cycles of refinement. The waters and other solvent molecules were added to the structures, and alternative conformations of residues were built using *COOT*. The refinement processes were rigorously monitored by assessing the reduction in *R_free_* values and ensuring that the overall stereochemical quality improved consistently throughout. The final refinement statistics, including completeness, *R_work_, R_free_* and stereochemical parameters for all structures presented in this study, are reported in Table 1.

## Acknowledgements

We thank Dr. Ravindra Makde, Dr. Ashwani Kumar and Dr. Biplab Ghosh at the PX-BL21 beamline (BARC) at Indus-2, RRCAT, Indore, India for their support in diffraction data collection. We acknowledge the Protein Crystallography Facility at IIT Bombay. We also thank Chinmay K. Kamale and Barnava Banerjee for proof reading the manuscript. The research was supported by research funding to PB from Department of Biotechnology, Government of India (grant number: BT/PR41982/PBD/26/822/2021).

## Author contributions

ABS, RKB and PB conceived the project. ABS, RKB collected the X-ray diffraction data. ABS, PB processed and refined the data and build model. ABS, RKB and MP designed and performed the experiments. ABS and PB wrote the manuscript with inputs from RKB, MP and SN

