## Supplementary Materials for "Rewiring Catalytic Craters: A Path for Engineering β-Glucosidases to Improve Glucose Tolerance"

*To whom correspondence should be addressed:

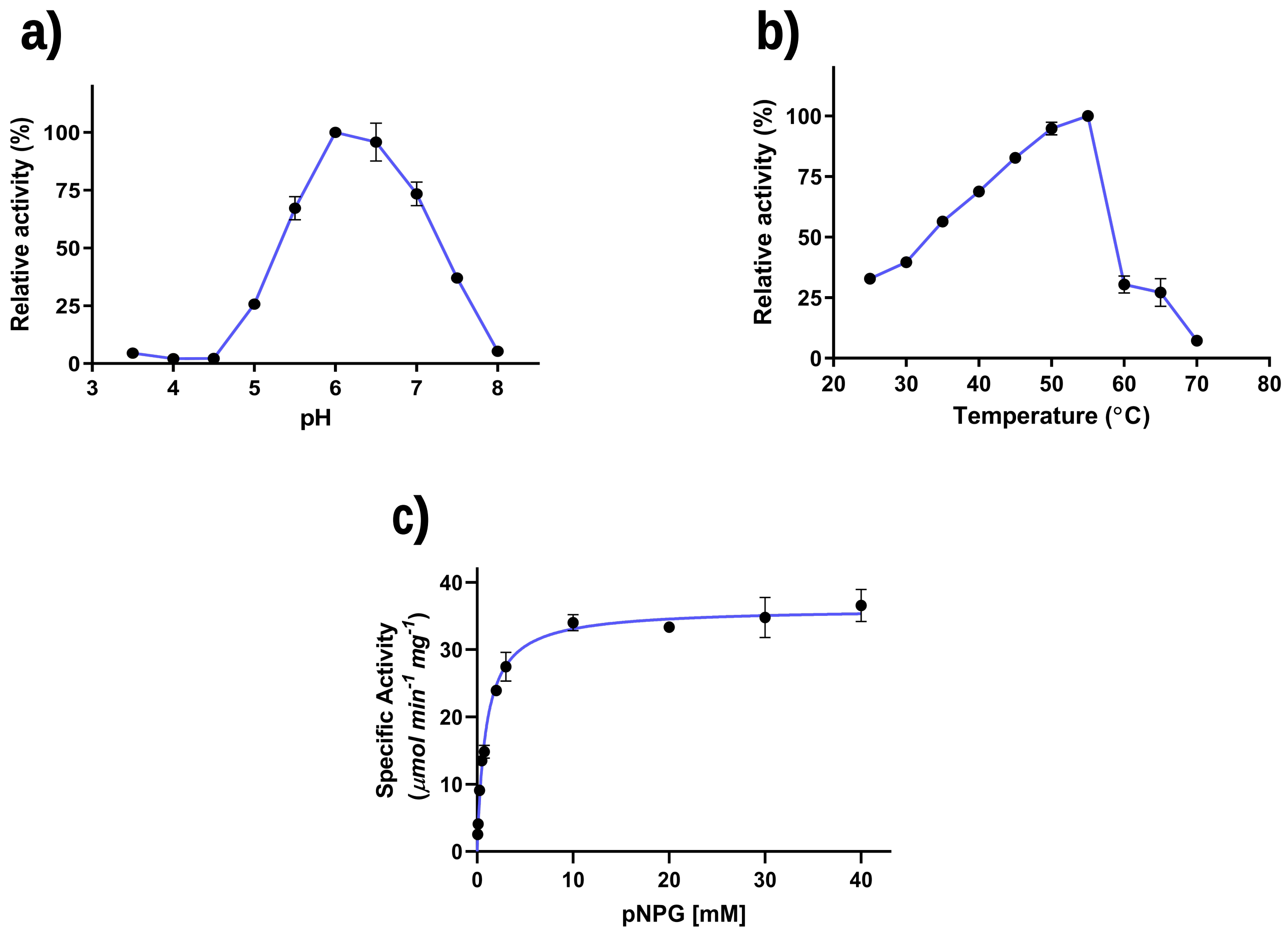


**Figure S1: a)** Graph illustrates the relative activity of pNPG hydrolysis pH profile for native UnBGl1, data points are shown as dark solid circle connected with purple line. Maximum activity shown at pH 6.0 and considered as 100 % activity. **b)** Graph illustrates the relative activity of pNPG hydrolysis for native UnBGl1 vs temperature profile, data points are shown as dark solid circle connected with purple line. Maximum activity shown at 55 °C is considered as 100 % activity. **c)** Kinetic characterization of native UnBGl1 with pNPG, showing the substrate saturation for determining the kinetic parameters. All experiments were performed in triplicates (n = 3) with error bar representing ± SEM.

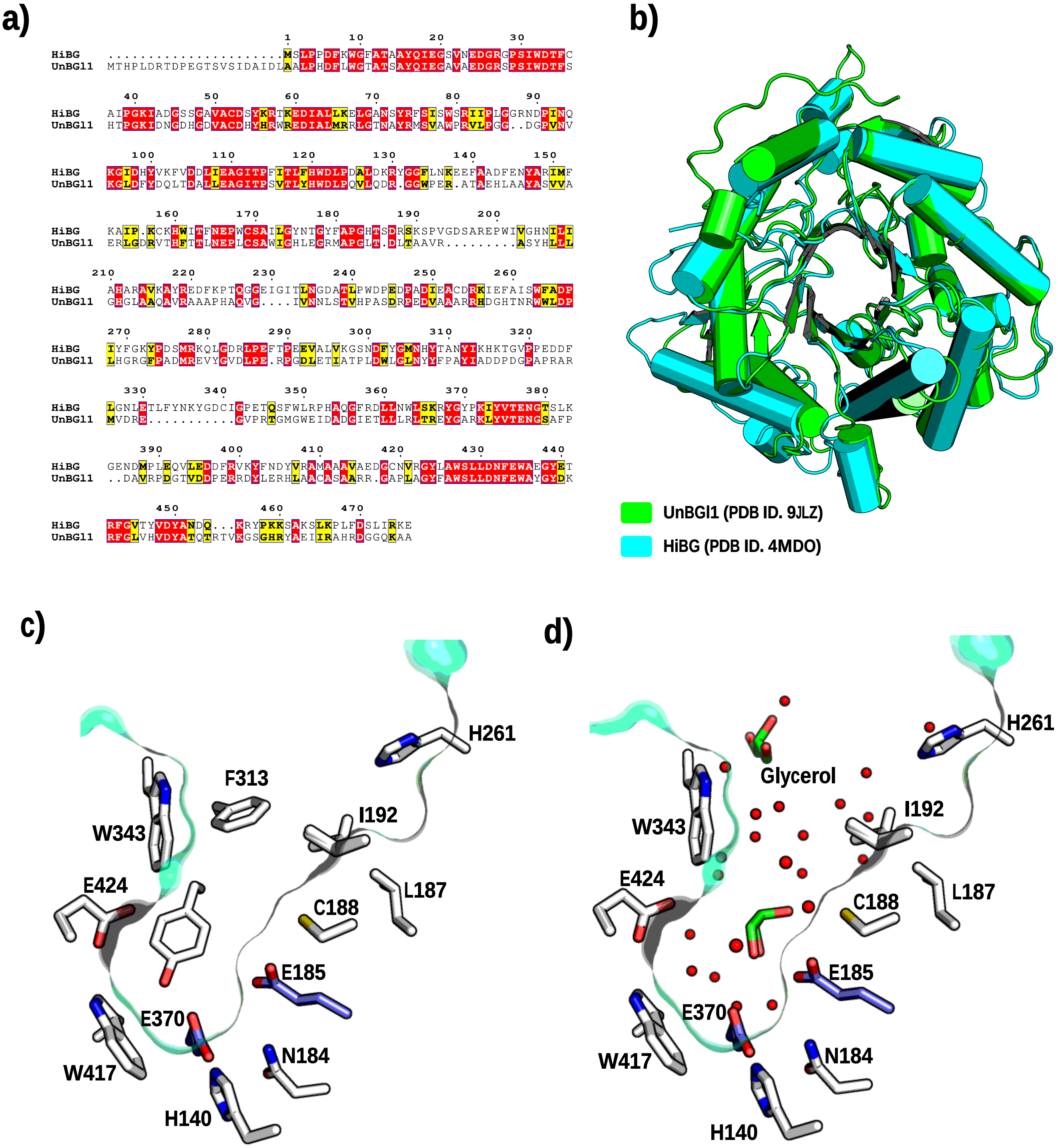


**Figure S2: a)** Sequence alignment between well characterized HiBG (β-glucosidase from *Humicola insolens* and UnBGl1 β-glucosidase enzyme performed using ESPript 3.0, highlighting the identical residues with red colour. **b)** Structural alignment of native UnBGl1 with HiBG shown in cartoon representation. **c)** UnBGl1 native crater shown in greencyan surface representation. Catalytic glutamates are shown in purple stick and all gatekeeper residues are shown in white sticks. **d)** UnBGl1 native crater shown in greencyan surface representation. Water molecules in the crater are represented as red spheres, glycerol present in the crater is shown in green sticks and gatekeeper residues are shown in white stick.

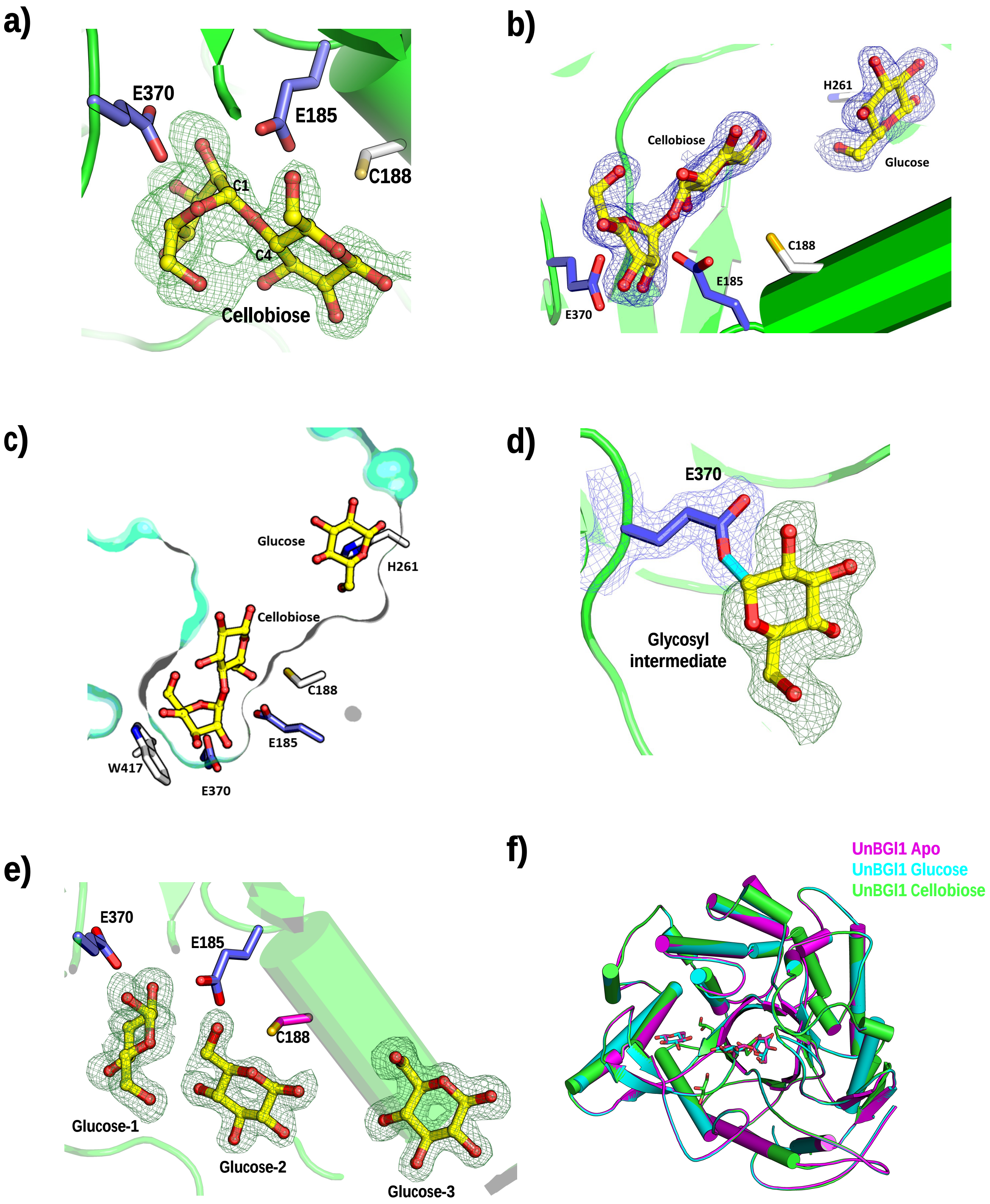
a)
E370
E185
C188
C1
C4
Cellobiose
b)
H261
Cellobiose
Glucose
C188
E185
E370
c)
Glucose
H261
Cellobiose
C188
E185
W417
E370
d)
E370
Glycosyl
intermediate
e)
E370
E185
C188
Glucose-1
Glucose-2
Glucose-3
f)
UnBGl1 Apo
UnBGl1 Glucose
UnBGl1 Cellobiose

**Figure S3: a)** Then $F_o$- $F_c$ omit map contoured at 3 σ level representing the electron density (forest green mesh) of cellobiose in near catalytic glutamates shown in purple sticks. **b)** The 2$F_o$ - $F_c$ map contoured at 1 σ level representing the electron density (blue mesh) of cellobiose near the catalytic glutamates (purple stick) and a glucose near H261$^{st}$ residue. **c)** Presence of cellobiose and glucose (yellow-ball and stick representation) shown in the crater of native UnBGl1. Crater is shown as greencyan surface. **d)** The 2$F_o$ - $F_c$ map contoured at 1 σ level representing the electron density (blue mesh) of catalytic glutamate (E370) (purple stick) forming a covalent bond (cyan) with a glucose molecule (yellow) shown in $F_o$ -$F_c$ map (forest green mesh) contoured at 3 σ level. **e)** The $F_o$ -$F_c$ omit map contoured 3 σ level representing the electron density (forest green mesh) three glucose molecules. **f)** Cartoon representation of structural alignment of native UnBGl1 in apo (magenta), glucose complexed (cyan) and cellobiose complexed (green) form representing no major structural difference between all the structures. Ligands in structures are shown in stick representation.

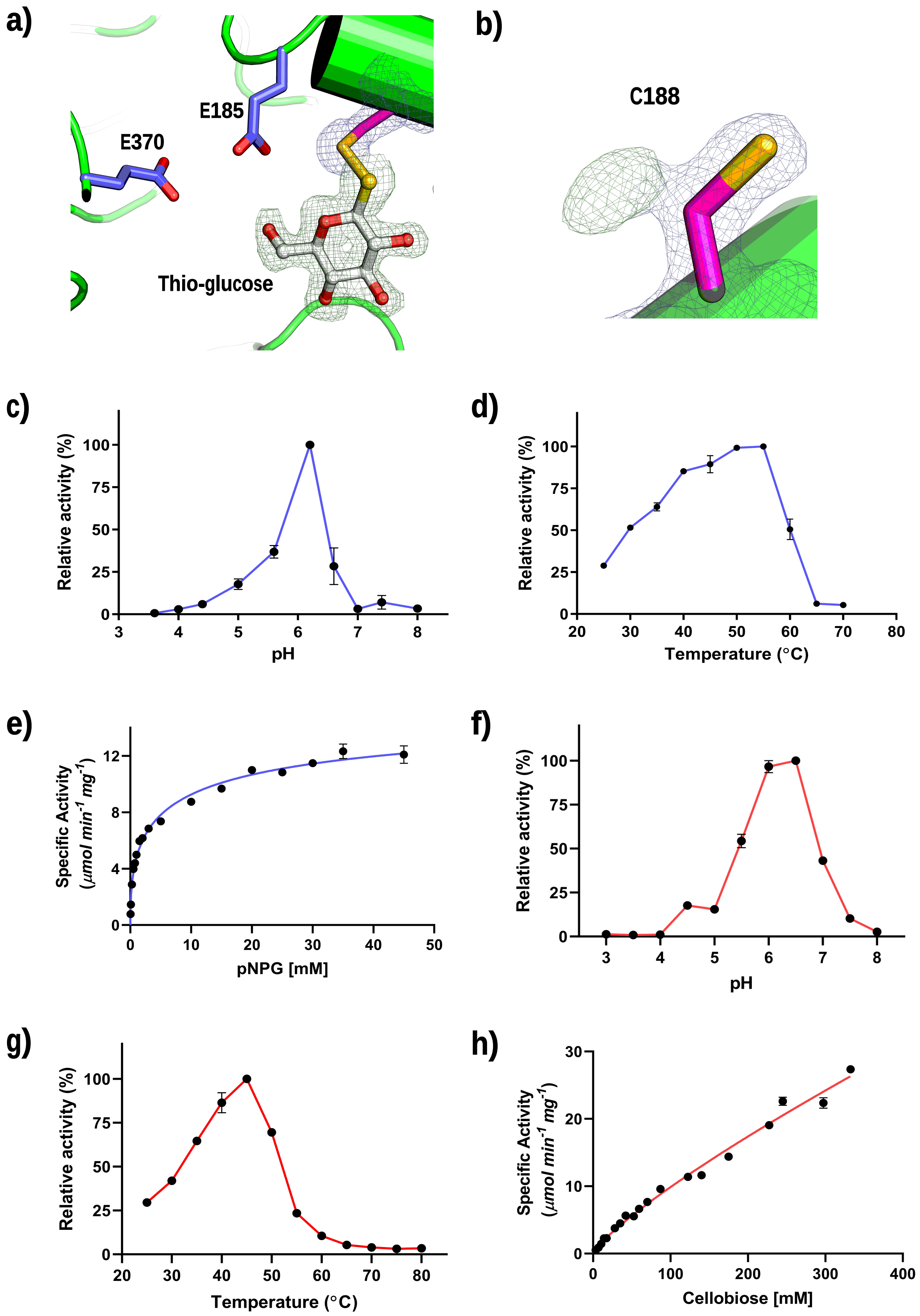

a)
E185
E370
Thio-glucose
b)
C188
c)
Relative activity (%)
pH
d)
Relative activity (%)
Temperature (°C)
e)
Specific Activity
(µmol min-1 mg-1)
pNPG [mM]
f)
Relative activity (%)
pH
g)
Relative activity (%)
Temperature (°C)
h)
Specific Activity
(µmol min-1 mg-1)
Cellobiose [mM]

**Figure S4: a)** The $2F_o - F_c$ map contoured at 1 σ level representing the electron density (slate mesh) of Cys188 (magenta stick) forming a disufide bond (golden color) with a thio-glucose molecule (white) shown in $F_o - F_c$ map (forest green mesh) contoured at 3 σ level. Catalytic glutamates are shown in purple stick. **b)** $2F_o - F_c$ map contoured at 1 σ level representing the electron density (slate mesh) of Cys188 (magenta stick) and $F_o - F_c$ map contoured at 3 σ level representing the extra electron density (forest green mesh) near cysteine in UnBGl1_C188V variant proving the presence of valine at 188$^{th}$ position. **c)** pH profile of UnBGl1_C188V for pNPG hydrolysis indicating optimum pH of 6.0, data points are shown as dark solid circle connected with purple line. **d)** Temperature profile of UnBGl1_C188V for pNPG hydrolysis indicating optimum temperature of 55 °C, data points are shown as dark solid circle connected with purple line. **e)** Kinetics profile of UnBGl1_C188V showing the substrate saturation for pNPG hydrolysis. **f)** pH profile of UnBGl1_C188V against hydrolysis of cellobiose at different pH, representing highest activity at pH 6.5. **g)** Temperature profile of UnBGl1_C188V for cellobiose hydrolysis indicating optimum temperature of 45 °C, data points are shown as dark solid circle connected with red line. **h) C**ellobiose vs the specific activity profile illustrating that UnBGl1_C188V requires higher concentration of cellobiose for saturation hence kinetic properties cannot be determined. Biochemical and kinetic assays were performed in triplicates (n = 3) with error bar representing ± SEM.

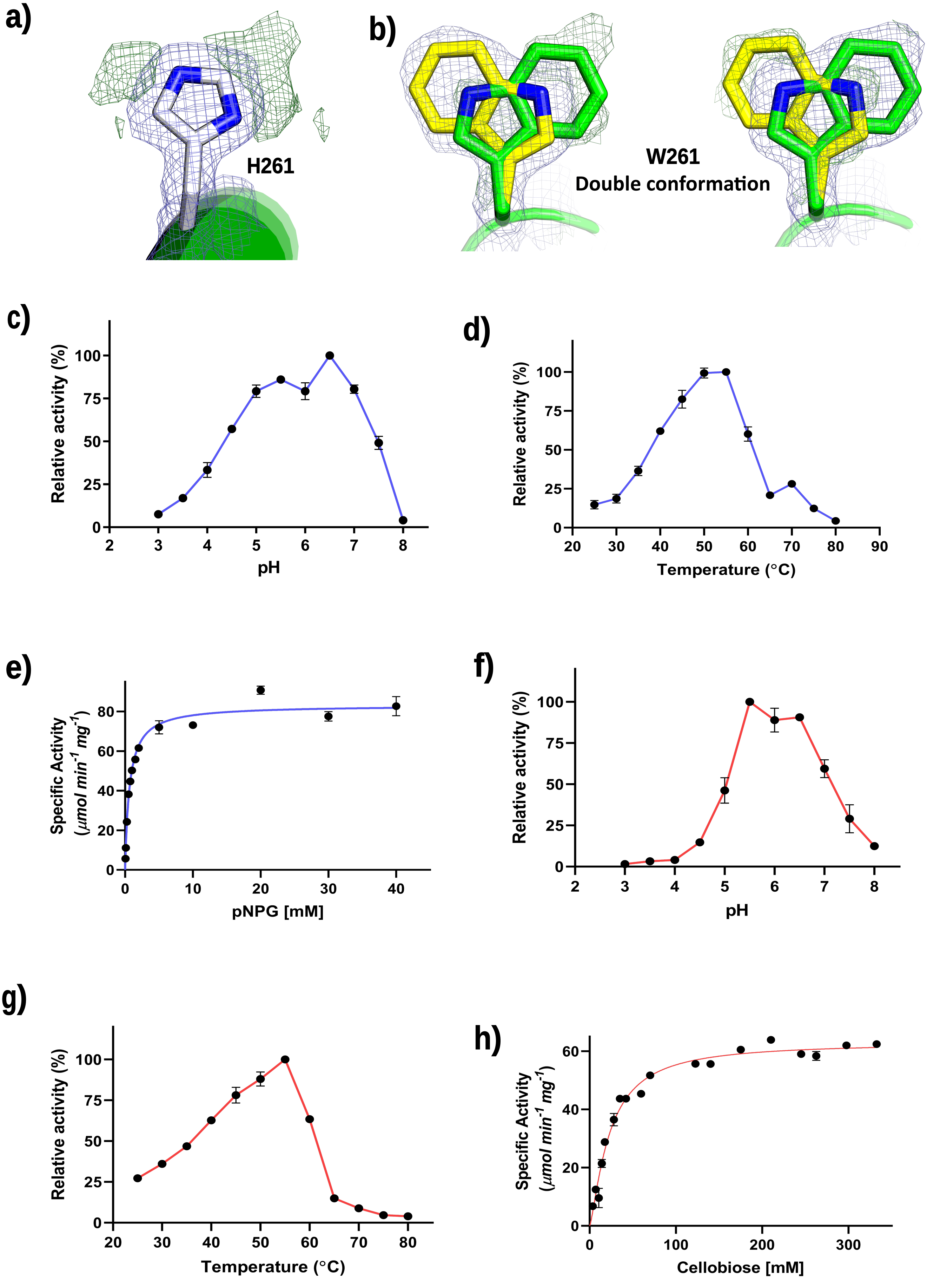
a)
H261
b)
W261
Double conformation
c)
Relative activity (%)
pH
d)
Relative activity (%)
Temperature (°C)
e)
Specific Activity
(µmol min-1 mg-1)
pNPG [mM]
f)
Relative activity (%)
pH
g)
Relative activity (%)
Temperature (°C)
h)
Specific Activity
(µmol min-1 mg-1)
Cellobiose [mM]

**Figure S5**: **a)** 2$F_o$ - $F_c$ map contoured at 1 σ level representing the electron density (slate mesh) of His261 (white stick). $F_o$- $F_c$ map contoured at 3 σ level representing extra electron density (forest green mesh) near His261 for tryptophan. **b)** $F_o$ - $F_c$ map (forest green mesh) contoured at 3 σ level representing alternate conformations of W261. **c)** pH profile of UnBGl1_H261W for pNPG hydrolysis indicating maximum activity at pH of 6.5, data points are shown as dark solid circle connected with purple line. **d)** Temperature profile of UnBGl1_H261W for pNPG hydrolysis indicating optimum temperature of 55 °C, data points are shown as dark solid circle connected with purple line. **e)** Kinetics profile of UnBGl1_H261W showing the substrate saturation for pNPG hydrolysis. **f)** pH profile of UnBGl1_H261W against hydrolysis of cellobiose at different pH, representing highest activity at pH 5.5, data points are shown as dark solid circle connected with red line. **g)** Temperature profile of UnBGl1_H261W for cellobiose hydrolysis indicating optimum temperature of 55 °C, data points are shown as dark solid circle connected with red line. **h)** Kinetic profile of UnBGl1_H261W for cellobiose hydrolysis. Biochemical and kinetic assays were performed in triplicates (n = 3) with error bar representing ± SEM.

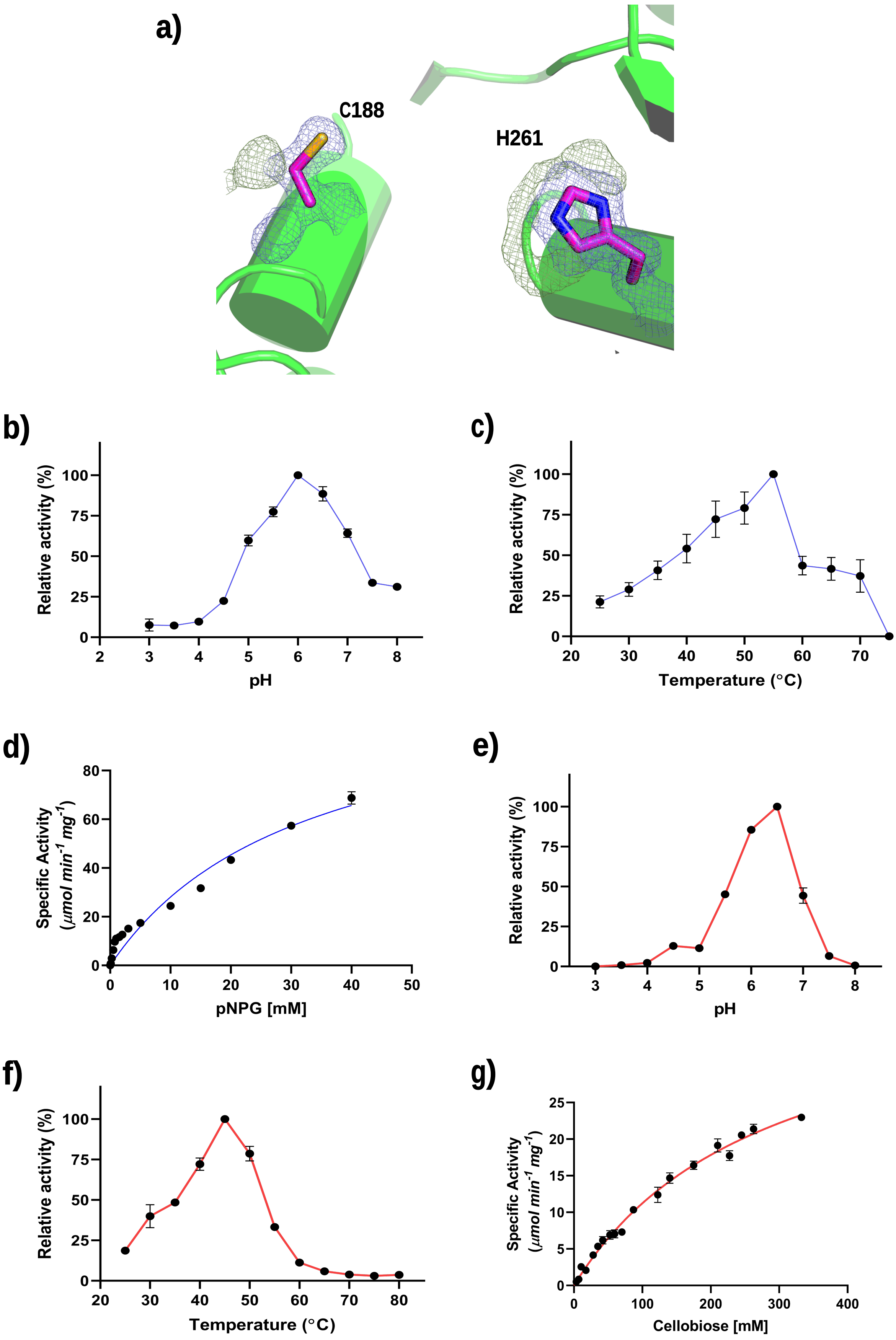
a)
C188
H261
b)
Relative activity (%)
100
75
50
25
0
2
3
4
5
6
7
8
pH
c)
Relative activity (%)
100
75
50
25
0
20
30
40
50
60
70
Temperature (°C)
d)
Specific Activity
(µmol min-1 mg-1)
80
60
40
20
0
0
10
20
30
40
50
pNPG [mM]
e)
Relative activity (%)
100
75
50
25
0
3
4
5
6
7
8
pH
f)
Relative activity (%)
100
75
50
25
0
20
30
40
50
60
70
80
Temperature (°C)
g)
Specific Activity
(µmol min-1 mg-1)
25
20
15
10
5
0
0
100
200
300
400
Cellobiose [mM]

**Figure S6: a)** The $2F_o - F_c$ map contoured at 1 σ level representing the electron density (slate mesh) of Cys188 and His261 (magenta stick). $F_o - F_c$ map contoured at 3 σ level representing extra electron density (forest green mesh) near Cys188 and His261 due presence of valine and tryptophan. **b)** Relative activity of pNPG hydrolysis by UnBGl1_C188V_H261W at different pH, data points are shown as dark solid circle connected with purple line. Maximum activity shown at pH 6.0 and considered as 100 % activity. **c)** Temperature profile of UnBGl1_C188V_H261W for hydrolysis of pNPG, showing maximum activity at 55 °C. **d)** Kinetic profile of UnBGl1 C188V_H261W for pNPG hydrolysis. **e)** pH profile of UnBGl1_C188V_H261W against hydrolysis of cellobiose at different pH, representing highest activity at pH 6.5. **f)** Temperature profile of UnBGl1_C188V_H261W for hydrolysis of cellobiose at different temperature, representing highest activity at 45 °C. **g)** Cellobiose vs the specific activity profile illustrating that UnBGl1_C188V_H261W requires higher concentration of cellobiose for saturation hence kinetic properties can not be determined. Biochemical and kinetic assays were performed in triplicates (n = 3) with error bar representing ± SEM.

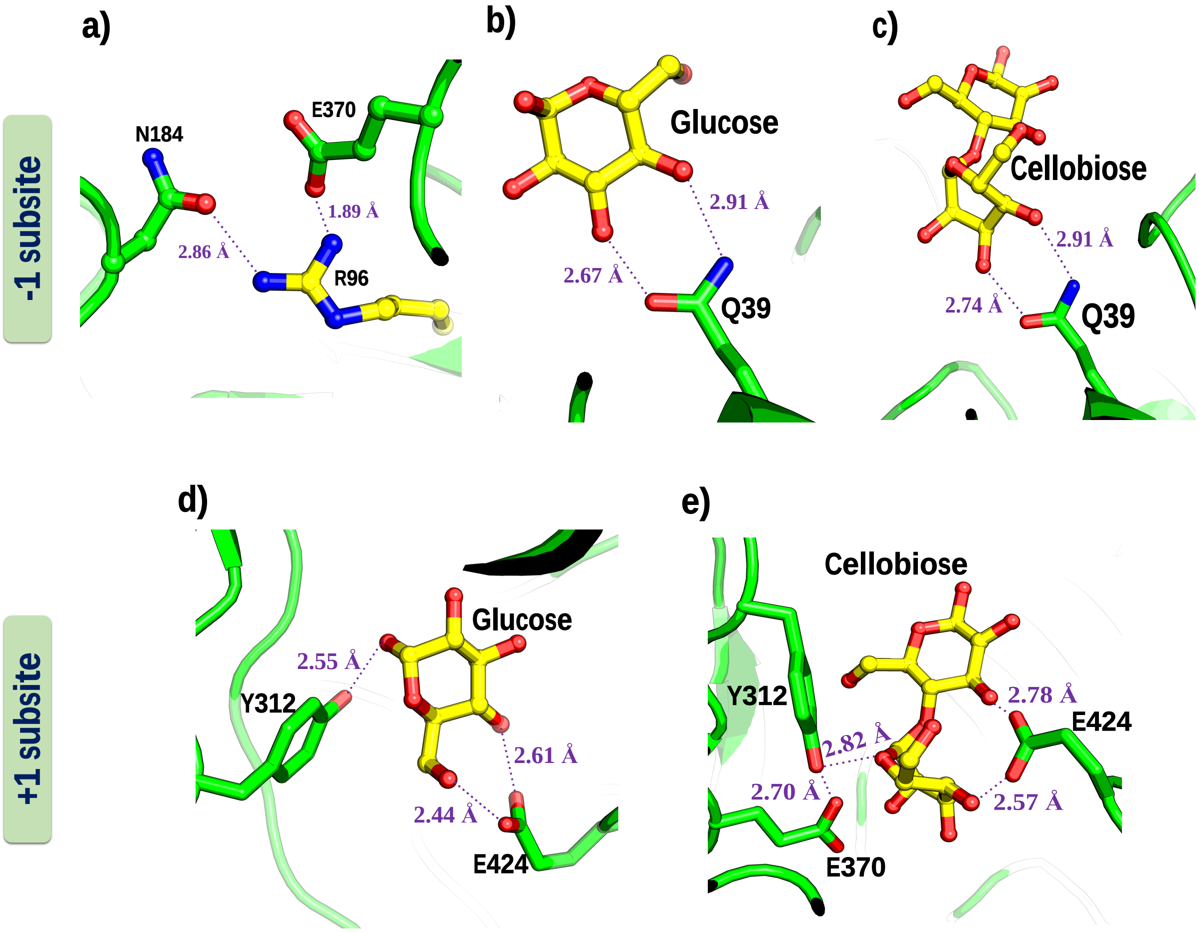


**Figure S7:** Glucose and cellobiose are shown in ball and stick presentation in yellow colour, residues interacting with the glucose and cellobiose are shown in green sticks, distances are shown in purple dash line **a)** NH1 and NH2 atom of R96 forming salt bridge interaction with OE2 of E370 and hydrogen bonding with OD1 of N184 respectively. **b)** NE2 and OE1 atoms of Q39 are forming hydrogen bond with O4 and O3 atom of glucose respectively. **c)** NE2 and OE1 atoms of Q39 are forming hydrogen bond with O8 and O7 atom of cellobiose respectively. **d)** OE1 and OE2 atoms of E424 forming hydrogen bonds with O4 and O6 atom of glucose respectively. OH atom of Y312 is present at hydrogen bonding distance from the O1 atom of glucose. **e)** OE1 and OE2 atoms of E424 forming hydrogen bonds with O8 and O4 atom of cellobiose respectively. OH atom of Y312 is present at hydrogen bonding distance from the O2 atom of cellobiose.

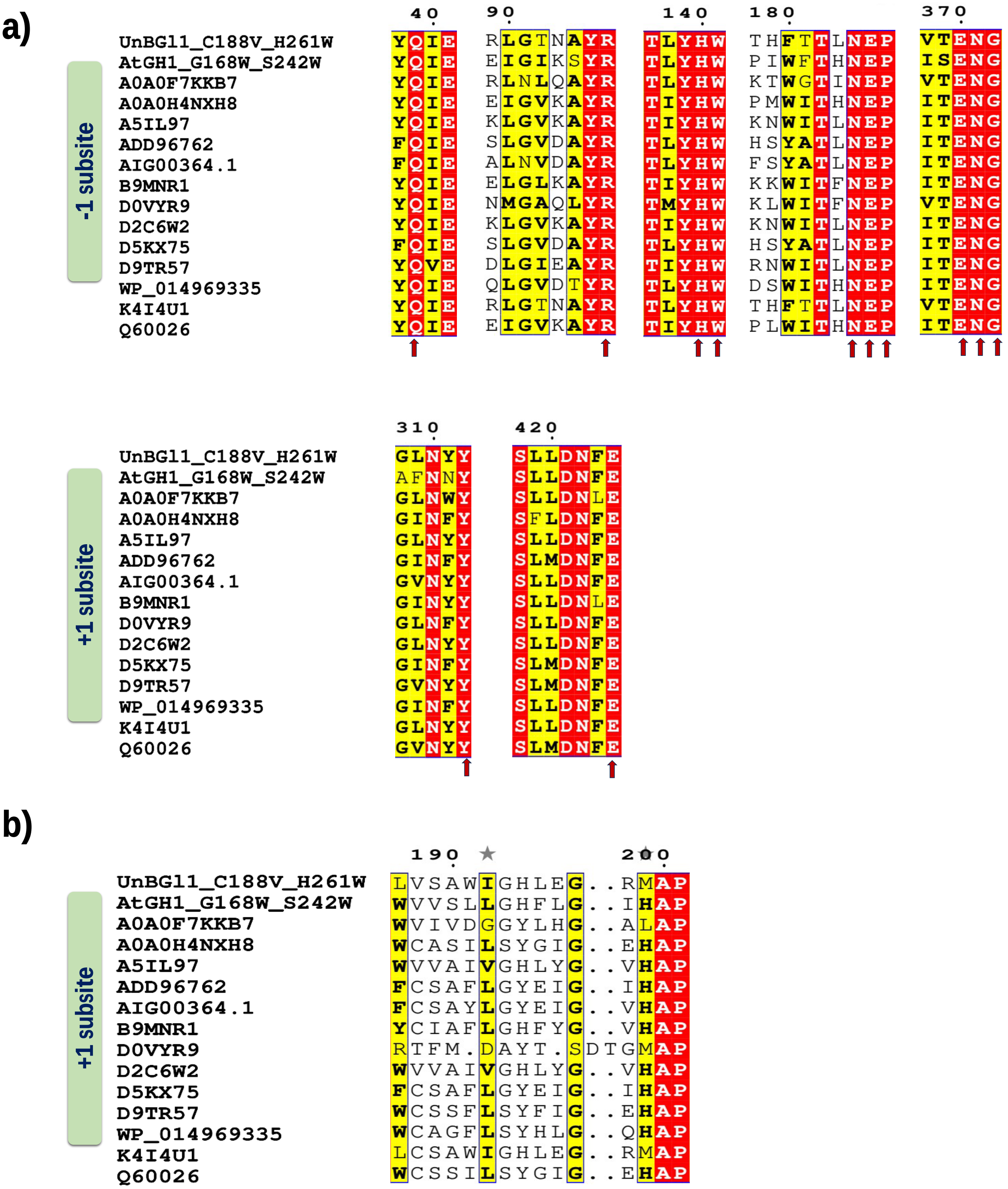
a)
-1 subsite
40 90 140 180 370
UnBGl1_C188V_H261W YQIE RLGTNAYR TLYHW THFTTLNEP VTENG
AtGH1_G168W_S242W YQIE EIGIKSYR TLYHW PIWFTHNEP ISENG
A0A0F7KKB7 YQIE RLNLQAYR TLYHW KTWGTINEP VTENG
A0A0H4NXH8 YQIE EIGVKAYR TIYHW PMWITHNEP ITENG
A5IL97 YQIE KLGVKAYR TIYHW KNWITLNEP ITENG
ADD96762 FQIE SLGVDAYR TLYHW HSYATLNEP ITENG
AIG00364.1 FQIE ALNVDAYR TLYHW FSYATLNEP ITENG
B9MNR1 YQIE ELGLKAYR TIYHW KKWITFNEP ITENG
D0VYR9 YQIE NMGAQLYR TMYHW KLWITFNEP VTENG
D2C6W2 YQIE KLGVKAYR TIYHW KNWITLNEP ITENG
D5KX75 FQIE SLGVDAYR TLYHW HSYATLNEP ITENG
D9TR57 YQVE DLGIEAYR TIYHW RNWITLNEP ITENG
WP_014969335 YQIE QLGVDTYR TLYHW DSWITHNEP ITENG
K4I4U1 YQIE RLGTNAYR TLYHW THFTTLNEP VTENG
Q60026 YQIE EIGVKAYR TIYHW PLWITHNEP ITENG
+1 subsite
310 420
UnBGl1_C188V_H261W GLNYY SLLDNFE
AtGH1_G168W_S242W AFNNY SLLDNFE
A0A0F7KKB7 GLNWY SLLDNLE
A0A0H4NXH8 GINFY SFLDNFE
A5IL97 GLNYY SLLDNFE
ADD96762 GINFY SLMDNFE
AIG00364.1 GVNYY SLLDNFE
B9MNR1 GINYY SLLDNLE
D0VYR9 GLNFY SLLDNFE
D2C6W2 GLNYY SLLDNFE
D5KX75 GINFY SLMDNFE
D9TR57 GVNYY SLMDNFE
WP_014969335 GINFY SLLDNFE
K4I4U1 GLNYY SLLDNFE
Q60026 GVNYY SLMDNFE
b)
+1 subsite
190 200
UnBGl1_C188V_H261W LVSAWIGHLEG..RMAP
AtGH1_G168W_S242W WVVSLLGHFLG..IHAP
A0A0F7KKB7 WVIVDGGYLHG..ALAP
A0A0H4NXH8 WCASILSYGIG..EHAP
A5IL97 WVVAIVGHLYG..VHAP
ADD96762 FCSAFLGYEIG..IHAP
AIG00364.1 FCSAYLGYEIG..VHAP
B9MNR1 YCIAFLGHFYG..VHAP
D0VYR9 RTFM.DAYT.SDTGMAP
D2C6W2 WVVAIVGHLYG..VHAP
D5KX75 FCSAFLGYEIG..IHAP
D9TR57 WCSSFLSYFIG..EHAP
WP_014969335 WCAGFLSYHLG..QHAP
K4I4U1 LCSAWIGHLEG..RMAP
Q60026 WCSSILSYGIG..EHAP

**Figure 8: a)** Structure based multiple sequence alignment of glucose tolerant GH1 β-glucosidase [*Acetivibrio thermocellus* (AtGH1_G168W_S242W), Uncultured bacterium (A0A0F7KKB7), *Thermoanaerobacterium aotearoense* (A0A0H4NXH8), *Thermotoga petrophila* (A5IL97), Uncultured bacterium (ADD96762), *Alteromonas australica* (AIG00364.1), *Caldicellulosiruptor bescii* (B9MNR1), *Nasutitermes takasagoensis* (D0VYR9), *Thermotoga naphthophila* (D2C6W2), Uncultured bacterium (D5KX75), *Thermoanaerobacterium thermosaccharolyticum* (D9TR57), *Exiguobacterium antarcticum* (WP_10496335), Uncultured bacterium (K4I4U1), *Thermoanaerobacter brockii* (Q60026)] with UnBGl1_C188V_H261W variant representing the conserved residues at -1 and +1 subsite that should not be modified for increasing glucose tolerance. Red arrows represent site of residues. **b)** Structure based multiple sequence alignment of glucose tolerant GH1 β-glucosidase with UnBGl1_C188V_H261W variant representing the conserved residues at +1 subsite that can be modified for increasing glucose tolerance. Red arrows represent site of residues

**Table S1:** List of primers

| | |
|---|---|
| ***Cloning*** | |
| *NdeI* | 5´ CAAATACATATGGAGAACCTGTACTTTCAAGGATGACCCATCCGC 3´ |
| *XhoI* | 5´ CTAGCTCGAGTTATGCTGCTTTCTGACCACCATCACGATG 3´ |
| ***Mutagenesis*** | |
| C188V Forward | 5´ GAATGAACCGCTGGTTAGCGCATGGATTGGTC 3´ |
| C188V Reverse | 5´ CATGCGCTAACCAGCAGTTCATTCAGGGT 3´ |
| H261W Forward | 5´ GCAGCAGCACGTTGGGATGGTCAT 3´ |
| H261W Reverse | 5´ GATTGGTATGACCATCCCAACGACGTG 3´ |

**Table S2:** List of cryoprotectants used for different variants (apo and complexed forms) of UnBGl1 crystals

| **Protein crystals** | **Crystal state during diffraction** | **Cryoprotectant** |
|---|---|---|
| Native UnBGl1 | Apo | 30 % glycerol |
| | Glucose Complexed | 1 M glucose |
| | Cellobiose Complexed | 40% PEG 3350 containing 250 mM cellobiose prepared in crystallization buffer |
| | Thio-cellobiose | 25 % glycerol |
| | Glycosyl intermediate | 40% PEG3350 containing 150 mM of cellobiose prepared in crystallization buffer |
| UnBGl1_C188V | Apo | 30 % glycerol in crystallization buffer |
| | Glucose Complexed | 2.3 M glucose prepared in crystallization buffer |
| UnBGl1_H261W | Apo | 35% glycerol in crystallization buffer |
| | Glucose Complexed | 2.3 M glucose prepared in crystallization buffer |
| | Cellobiose Complexed | 20 % glycerol in 300 mM cellobiose prepared in crystallization buffer |
| UnBGl1_C188V_H261W | Apo | 35% glycerol in crystallization buffer |
| | Glucose Complexed | 2 M glucose prepared in crystallization buffer having 20% glycerol |
